# Genomic basis of parallel adaptation varies with divergence in *Arabidopsis* and its relatives

**DOI:** 10.1101/2020.03.24.005397

**Authors:** Magdalena Bohutínská, Jakub Vlček, Sivan Yair, Benjamin Laenen, Veronika Konečná, Marco Fracassetti, Tanja Slotte, Filip Kolář

## Abstract

Parallel adaptation provides valuable insight into the predictability of evolutionary change through replicated natural experiments. A steadily increasing number of studies have demonstrated genomic parallelism, yet the magnitude of this parallelism varies depending on whether populations, species or genera are compared. This led us to hypothesize that the magnitude of genomic parallelism scales with genetic divergence between lineages, but whether this is the case and the underlying evolutionary processes remain unknown. Here, we resequenced seven parallel lineages of two *Arabidopsis* species which repeatedly adapted to challenging alpine environments. By combining genome-wide divergence scans with model-based approaches we detected a suite of 151 genes that show parallel signatures of positive selection associated with alpine colonization, involved in response to cold, high radiation, short season, herbivores and pathogens. We complemented these parallel candidates with published gene lists from five additional alpine Brassicaceae and tested our hypothesis on a broad scale spanning ~ 0.02 to 18 million years of divergence. Indeed, we found quantitatively variable genomic parallelism whose extent significantly decreased with increasing divergence between the compared lineages. We further modeled parallel evolution over the *Arabidopsis* candidate genes and showed that a decreasing probability of repeated selection of the same standing or introgressed alleles drives the observed pattern of divergence-dependent parallelism. We therefore conclude that genetic divergence between populations, species and genera, affecting the pool of shared variants, is an important factor in the predictability of genome evolution.

**Significance statement:** Repeated evolution tends to be more predictable. The impressive spectrum of recent reports on genomic parallelism, however, revealed that the fraction of the genome that evolves in parallel largely varies, possibly reflecting different evolutionary scales investigated. Here, we demonstrate divergence-dependent parallelism using a comprehensive genome-wide dataset comprising 12 cases of parallel alpine adaptation and identify decreasing probability of adaptive re-use of genetic variation as the major underlying cause. This finding empirically demonstrates that evolutionary predictability is scale dependent and suggests that availability of pre-existing variation drives parallelism within and among populations and species. Altogether, our results inform the ongoing discussion about the (un)predictability of evolution, relevant for applications in pest control, nature conservation, or the evolution of pathogen resistance.

## Introduction

Evolution is driven by a complex interplay of deterministic and stochastic forces whose relative importance is a matter of debate (1). Being largely a historical process, we have limited ability to experimentally test for the predictability of evolution in its full complexity, i.e., in natural environments (2). Distinct lineages that independently adapted to similar conditions by similar phenotype (termed ‘parallel’, considered synonymous to ‘convergent’ here) can provide invaluable insights into the issue (3, 4). An improved understanding of the probability of parallel evolution in nature may inform on constraints on evolutionary change and provide insights relevant for predicting the evolution of pathogens (5–7), pests (8, 9) or species in human-polluted environments (10, 11). Although the past few decades have seen an increasing body of work supporting the parallel emergence of traits by the same genes and even alleles, we know surprisingly little about what makes parallel evolution more likely and, by extension, what factors underlie evolutionary predictability (1, 12).

A wealth of literature describes the probability of ‘genetic’ parallelism, showing why certain genes are involved in parallel adaptation more often than others (13). There is theoretical and empirical evidence for the effect of pleiotropic constraints, availability of beneficial mutations or position in the regulatory network all having an impact on the degree of parallelism at the level of a single locus (3, 13–18). In contrast, we know little about causes underlying ‘genomic’ parallelism, i.e., what fraction of the genome is reused in adaptation and why. Individual case studies demonstrate large variation in genomic parallelism, ranging from absence of any parallelism (19), similarity in functional pathways but not genes (20, 21), reuse of a limited number of genes (22, 23) to abundant parallelism at both gene and functional levels (24, 25). Yet, there is little consensus about what determines variation in the degree of gene reuse (fraction of genes that repeatedly emerge as selection candidates) across investigated systems (1).

Divergence (the term used here to consistently describe both intra- and interspecific genetic differentiation) between the compared instances of parallelism appears as a promising candidate (14, 26, 27). Phenotype-oriented meta-analyses suggest that both phenotypic convergence (27) and genetic parallelism underlying phenotypic traits (14) decrease with increasing time to the common ancestor. Although a similar targeted multi-scale comparison is lacking at the genomic level, our brief review of published studies (29 cases, Dataset S1) suggests that also gene reuse tends to scale with divergence (Fig. 1a, Fig. S1). Moreover, allele reuse (repeated sweep of the same haplotype that is shared among populations either via gene flow or from standing genetic variation) frequently underlies parallel adaptation between closely related lineages (28–31), while parallelism from independent *de-novo* mutations at the same locus dominates between distantly related taxa (13). Similarly, previous studies reported a decreasing probability of hemiplasy (apparent convergence resulting from gene tree discordance) with divergence in phylogeny-based studies (32, 33). This suggests that the degree of allele reuse may be the primary factor underlying the hypothesized divergence-dependency of parallel genome evolution, possibly reflecting either genetic (weak hybridization barriers, widespread ancestral polymorphism between closely related lineages (34) or ecological reasons (lower niche differentiation and geographical proximity (35, 36). However, the generally restricted focus of individual studies of genomic parallelism on a single level of divergence does not lend itself to a unified comparison across divergence scales. Although different ages of compared lineages affects a variety of evolutionary-ecological processes such as diversification rates, community structure or niche conservatism (36), the hypothesis that genomic parallelism scales with divergence has not yet been systematically tested and the underlying evolutionary processes remain poorly understood.

**Figure 1:**
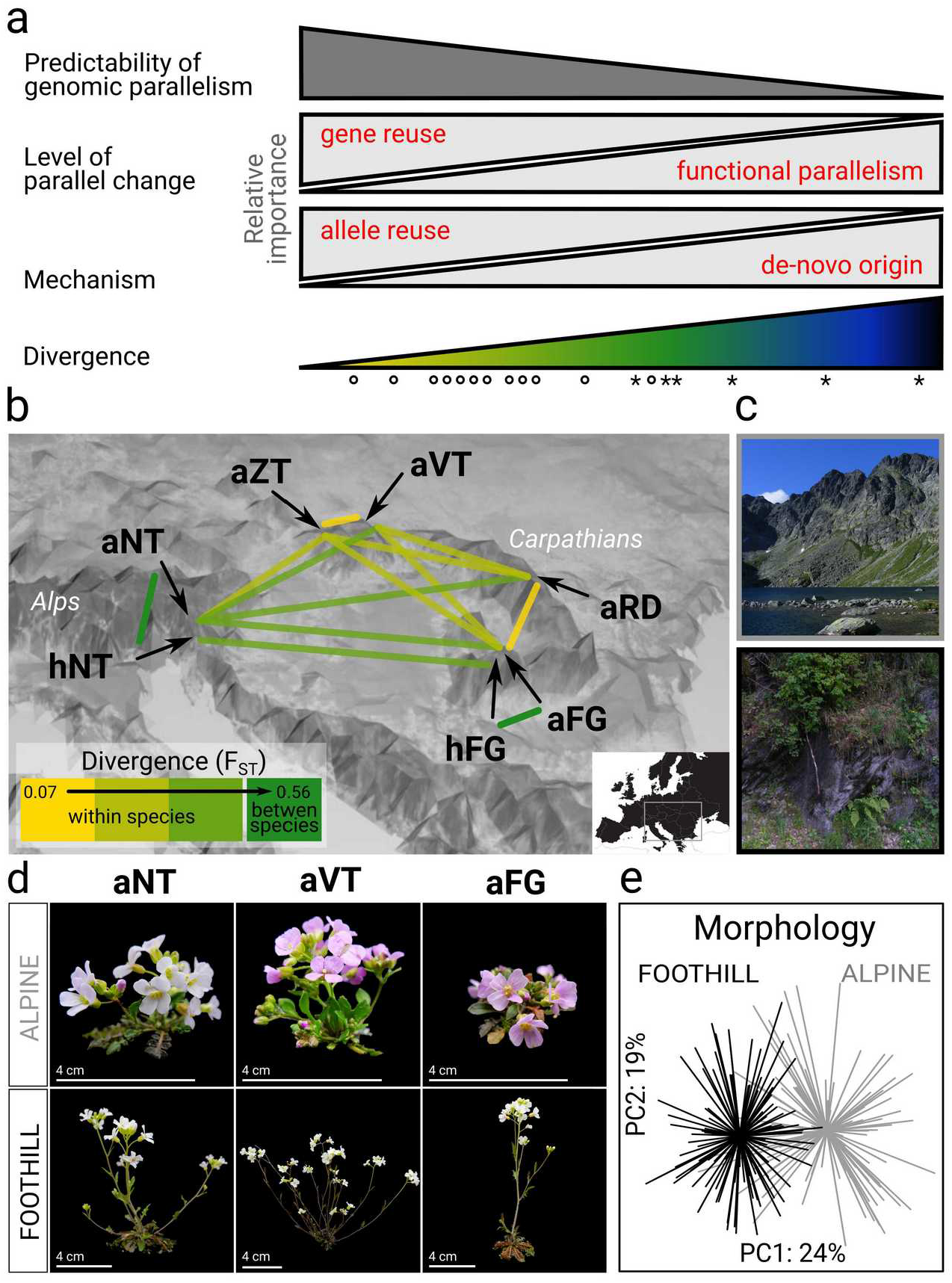
Hypotheses regarding relationships between genomic parallelism and divergence and the *Arabidopsis* system used to address these hypotheses. (a) Based on our literature review we propose that genetically closer lineages adapt to a similar challenge more frequently by gene reuse, sampling suitable variants from the shared pool (allele reuse), which makes their adaptive evolution more predictable. Color ramp symbolizes rising divergence between the lineages (~0.02-18 Mya in this study), the symbols denote different divergence levels tested here using resequenced genomes of 22 *Arabidopsis* populations (circles) and meta-analysis of candidates in Brassicaceae (asterisks). (b) Spatial arrangement of lineages of varying divergence (neutral F_ST_; bins only aid visualization, all tests were performed on a continuous scale) encompassing parallel alpine colonization within the two *Arabidopsis* outcrosses from central Europe: *A. arenosa* (diploid: aVT, autotetraploid: aNT, aZT, aRD and aFG) and *A. halleri* (diploid: hNT and hFG). Note that only two of the ten between-species pairs (dark green) are shown to aid visibility. The color scale corresponds to left part of a ramp used in a). (c) Photos of representative alpine and foothill habitat. (d) Representative phenotypes of originally foothill and alpine populations grown in common garden demonstrating phenotypic convergence. (e) Morphological differentiation among 223 *A. arenosa* individuals originating from foothill (black) and alpine (gray) populations from four regions after two generations in a common garden. Principal component analysis was run using 16 morphological traits taken from (44).

Here, we aimed to test this hypothesis and investigate whether allele reuse is a major factor underlying the relationship. We analyzed replicated instances of adaptation to a challenging alpine environment, spanning a range of divergence from populations to tribes within the plant family Brassicaceae (37–42) (Fig. 1a). First, we took advantage of a unique naturally multi-replicated setup in the plant model genus *Arabidopsis,* that was so far neglected from a genomic perspective (Fig. 1b). Two predominantly foothill-dwelling *Arabidopsis* outcrossers (*A. arenosa, A. halleri*) exhibit scattered, morphologically distinct alpine occurrences at rocky outcrops above the timberline (Fig. 1c). These alpine forms are separated from the widespread foothill population by a distribution gap spanning at least 500 m of elevation. Previous genetic and phenotypic investigations and follow-up analyses presented here showed that the scattered alpine forms of both species represent independent alpine colonization in each mountain range, followed by parallel phenotypic differentiation (Fig. 1d, e) (43, 44). Thus, we sequenced genomes from seven alpine and adjacent foothill population pairs, covering all European lineages encompassing the alpine ecotype. We discovered a suite of 151 genes from multiple functional pathways relevant to alpine stress, that were repeatedly differentiated between foothill and alpine populations. This points towards a polygenic, multi-factorial basis of parallel alpine adaptation.

We took advantage of this set of well-defined parallel selection candidates and tested whether the degree of gene reuse decreases with increasing divergence between the compared lineages (Fig. 1a). By extending our analysis to five additional alpine Brassicaceae species, we further tested whether there are limits to gene reuse above the species level. Finally, we inquired about possible underlying evolutionary process by estimating the extent of allele reuse using a designated modeling approach. Overall, our empirical analysis provides a novel perspective to the ongoing discussion about the variability in the reported magnitude of parallel genome evolution and identifies allele reuse as an important evolutionary process shaping the extent of genomic parallelism between populations, species and genera.

## Results

### Parallel alpine colonization by distinct lineages of *Arabidopsis*

We retrieved whole genome sequences from 11 alpine and 11 nearby foothill populations (174 individuals in total, seven to eight per population) covering all seven mountain regions with known occurrence of *Arabidopsis arenosa* or *A. halleri* alpine forms (a set of populations from one mountain region is further referred to as a ‘lineage’, Fig. 1b, Fig. S2, Table S1, S2). Within each species, population structure analyses based on genome-wide nearly-neutral four-fold degenerate (4d) SNPs demonstrated clear grouping according to lineage but not alpine environment, suggesting parallel alpine colonization of each mountain region by a distinct genetic lineage (Fig. S3, S4). This was in line with separation histories between diploid populations of *A. halleri* estimated in Relate (Fig. S5) and previous coalescent simulations on broader population sampling of *A. arenosa* (44). The only exception was the two spatially closest lineages of *A. arenosa* (aVT and aZT) for which alpine populations clustered together, keeping the corresponding foothill populations paraphyletic. Due to considerable pre-(spatial segregation) and post-zygotic (ploidy difference) barriers between the alpine populations from these two lineages (45) we left aZT and aVT as separate units in the following analyses for the sake of clarity (exclusion of this pair of lineages did not lead to qualitatively different results, Supplementary Text 1).

We observed a gradient of neutral differentiation among the seven lineages (quantified as average pairwise 4d-F_ST_ between foothill populations from each lineage, range 0.07 – 0.56, Table S3). These values correlated with absolute neutral divergence (4d-D_XY_, Pearson’s r = 0.89, p < 0.0001) and we further refer to them consistently as ‘divergence’. All populations showed high levels of 4d-nucleotide diversity (mean = 0.023, SD = 0.005), as expected for strict outcrossers and no remarkable deviation from neutrality (the range of 4d-Tajima’s D was −0.16 – 0.66, Supplementary Table 4). We found no signs of severe demographic change that would be associated with alpine colonization (similar 4d-nucleotide diversity and 4d-Tajima’s D of alpine and foothill populations; Wilcoxon rank test, p = 0.70 and 0.92, respectively, n = 22). Coalescent-based demographic inference further supported a no-bottleneck model even for the outlier population with the highest positive 4d-Tajima’s D (population LAC of aFG lineage, Fig. S6).

### Genomic basis of parallel alpine adaptation

Leveraging the seven natural replicates, we identified a set of genes showing signatures of parallel directional selection associated with alpine colonization. We used a conservative approach taking the intersection of F_ST_-based divergence scans designed to control for potential confounding signal of local selection within each ecotype (see Methods) and candidate detection under a Bayesian framework that accounts for neutral processes (BayPass) and identified 100 – 716 gene candidates in each of the seven lineages. Of these, we identified 196 gene candidates that were shared between at least two lineages and further tested whether they are consistent with parallel adaptation using neutral simulations in the Distinguishing Modes of Convergence (DMC) maximum composite likelihood framework (46) (see Methods for details). Out of the 196 shared gene candidates, we identified 151 genes showing significantly higher support for the parallel alpine selection model as compared to a neutral model assuming no selection in DMC (further referred to as ‘parallel gene candidates’). This set of genes contains an enrichment of differentiated nonsynonymous SNPs (Table S5) and we did not find any evidence that this was explained by weaker selective constraint compared to the rest of the genome (approximated by ratio of their nonsynonymous to synonymous diversity, Table S6, Fig. S7 and Supplementary Text 2). Further, the parallel gene candidates do not group together nor are they clustered in the regions of low recombination rates (Fig. S8, S9). Functional annotations of the parallel gene candidates using the TAIR database and associated publications (Dataset S2), protein-protein interaction database STRING (Fig. S10) and gene onthology (GO) enrichment analysis (Dataset S3) suggest a complex polygenic basis of alpine adaptation, involving multiple major functional categories, well-matching expectations for a response to a multifactorial environmental stress (Supplementary Text 3). Six of the physiological adaptations to alpine environment, encompassing both abiotic and biotic stress stand out (broadly following (47)), both in terms of number of associated parallel candidate genes and functional pathways (Fig. 2). We further discuss these putative alpine adaptations and their functional implications in Supplementary Text 3.

**Figure 2:**
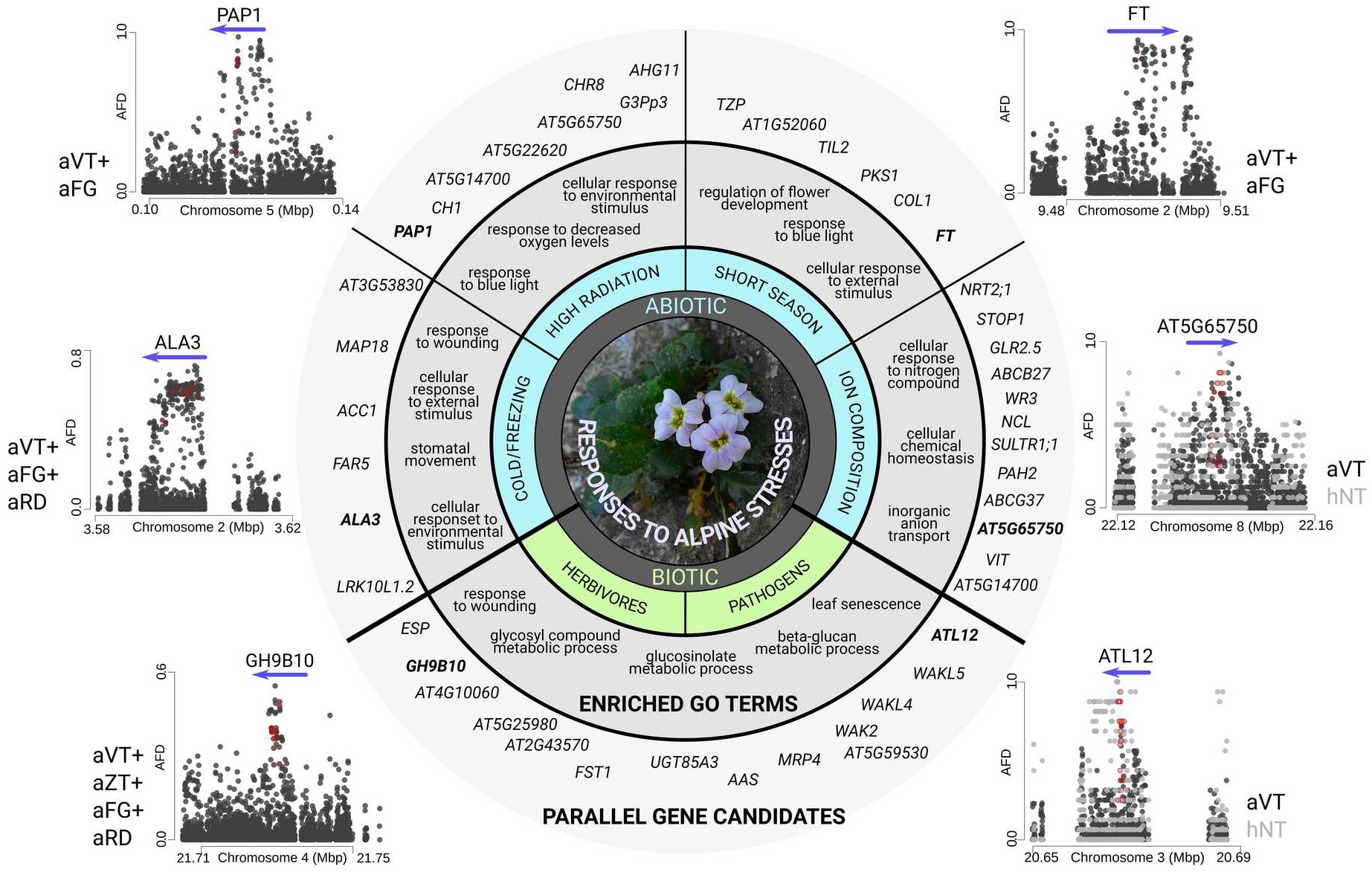
Physiological responses to alpine stresses in *Arabidopsis arenosa* and *A. halleri*, identified based on functional annotation of parallel gene candidates (circle) and signatures of parallel directional selection at the corresponding loci (surrounding dotplots). The circle scheme is based on the annotated list of 151 parallel gene candidates (Dataset S2) and corresponding enriched GO terms within the biological process category (Dataset S3). For more details on functional interpretations see Supplementary Text 3. Dotplots show allele frequency difference (AFD) at single-nucleotide polymorphisms between foothill and alpine populations summed over all lineages showing a parallel differentiation in a given gene (blue arrow). The lineage names are listed on the sides. Loci with two independently differentiated haplotypes likely representing independent *de-novo* mutations (AT5G65750 and ATL12) are represented by peaks of black and gray dots, corresponding with the two parallel lineages. Red circles highlight nonsynonymous variants.

### Ubiquitous gene and function-level parallelism and their relationship with divergence

Using the set of alpine gene candidates identified in *Arabidopsis* lineages, we quantified the degree of parallelism at the level of genes and gene functions (biological processes). We overlapped the seven lineage-specific candidate gene lists across all 21 pairwise combinations of the lineages and identified significant parallelism (non-random number of overlapping genes, p < 0.05, Fisher’s exact test, Fig. 3a), among 15 (71 %) lineage pairs (Table S7). Notably, the overlaps were significant for 10 out of 11 pairwise comparisons among the lineages within a species but only in five out of 10 pairwise comparisons across species (Dataset S4). We then annotated the functions of gene candidates using ‘biological process’ GO terms in each lineage, extracted only significantly enriched functions and again overlapped them across the seven lineages. Of these, we found significant overlaps (p < 0.05, Fisher’s exact test) among 17 (81 %) lineage pairs and the degree of overlap was similar within and across species (82 and 80 %, respectively, Fig. 3b, Table S7, Dataset S5).

**Figure 3:**
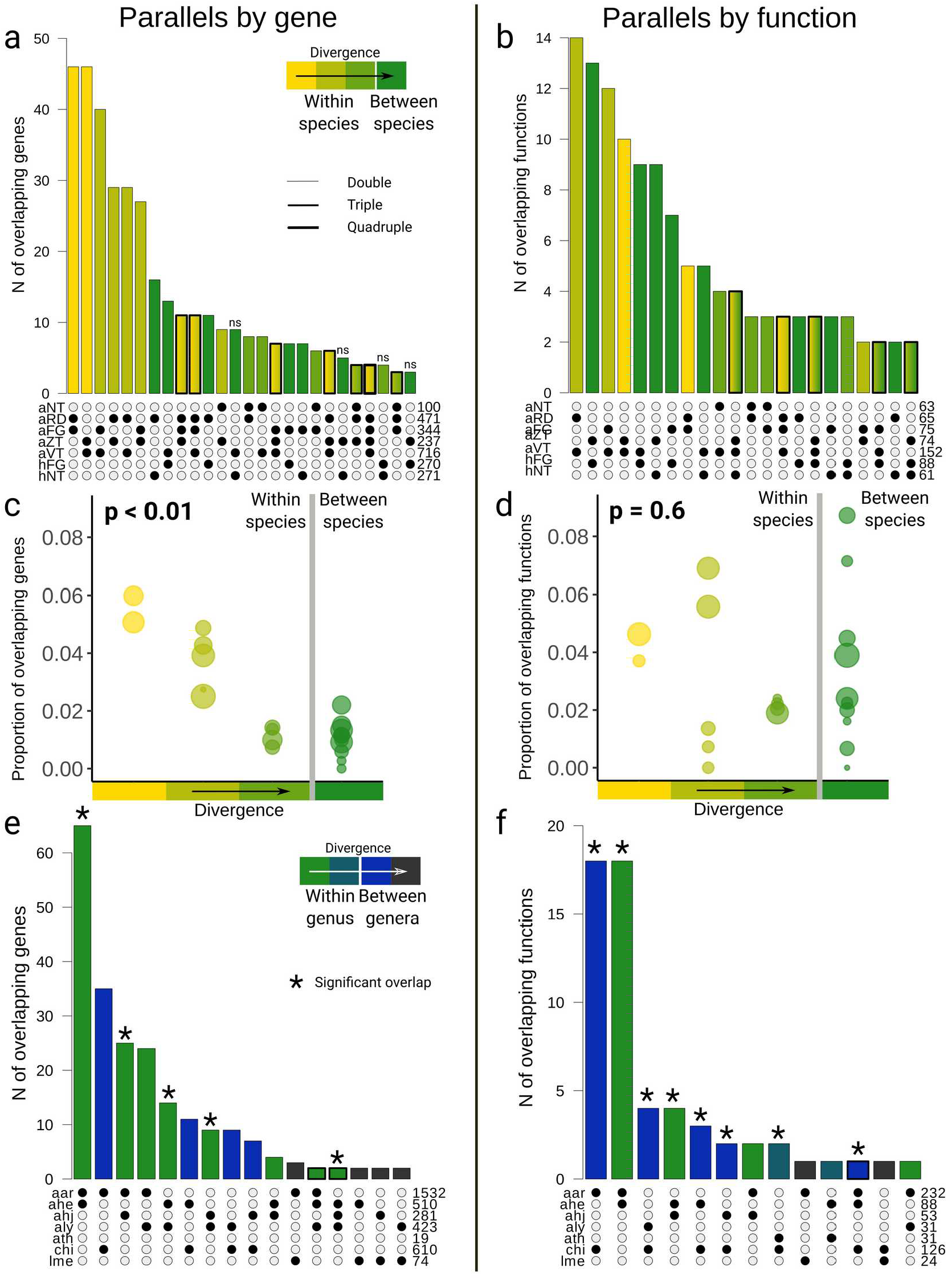
Variation in gene and function-level parallelism and their relationship with divergence in *Arabidopsis arenosa* and *A. halleri* (a-d) and across species from Brassicaceae family (e, f). (a, b) Number of overlapping candidate genes (a) and functions (b; enriched GO terms) for alpine adaptation colored by increasing divergence between the compared lineages. Only overlaps of > 2 genes and > 1 function are shown (for a complete overview see Datasets S4-S7). Numbers in the bottom-right corner of each panel show the total number of candidates in each lineage. Unless indicated (“ns”) the categories exhibited higher than random overlap of the candidates (p < 0.05, Fisher’s exact test). For lineage codes see Fig. 1b. Categories with overlap over more than two lineages are framed in bold and filled by a gradient. (c, d) Proportions of parallel genes (c; gene reuse) and functions (d) among all candidates identified within each pair of lineages (dot) binned into categories of increasing divergence (bins correspond to Fig. 1b) with significance levels inferred by Mantel test over continuous divergence scale (see the text). Size of the dot corresponds to the number of parallel items. (e, f) Same as (a, b) but for species from Brassicaceae family, spanning higher divergence levels. Categories indicated by an asterisk exhibited higher than random overlap of the candidates (p < 0.05, Fisher’s exact test). Codes: aar: our data on *Arabidopsis arenosa* ahe: our data on *A. halleri* combined with *A. halleri* candidates from Swiss Alps (38), ahj: *Arabidopsis halleri* subsp. *gemmifera* from Japan (37), aly: *Arabidopsis lyrata* from Northern Europe (39), ath *: Arabidopsis thaliana* from Alps (42), chi: *Crucihimalaya himalaica* (41), lme: *Lepidium meyenii* (40).

Then, we quantified the degree of parallelism for each pair of *Arabidopsis* lineages as the proportion of overlapping gene and function candidates out of all candidates identified for these two lineages. The degree of parallelism was significantly higher at the function level (mean proportion of parallel genes and functions across all pairwise comparisons = 0.045 and 0.063, respectively, D = 437.14, df = 1, p < 0.0001, GLM with binomial errors). Importantly, the degree of parallelism at the gene-level (i.e., gene reuse) significantly decreased with increasing divergence between the lineages (negative relationship between Jaccard’s similarity in candidate gene identity among lineages and 4d-F_ST_; Mantel rM = −0.71, p = 0.001, 999 permutations, Fig. 3c). In contrast, the degree of parallelism by function did not correlate with divergence (rM = 0.06, p = 0.6, 999 permutations, n = 21, Fig. 3d).

We further tested whether the relationship between the degree of parallelism and divergence persists at deeper phylogenetic scales by complementing our data with candidate gene lists from six genome-wide studies of alpine adaptation from the Brassicaceae family (37–42) (involving five species diverging 0.5 – 18 millions of years ago (48, 49), SI Appendix, Methods; Table S8, S9). While we still found significant parallelism both at the level of candidate genes and functions (Figure 3e, f, Dataset S6, S7), their relationship with divergence was non-significant (Mantel rM = −0.52 / −0.22, for genes / functions respectively, p = 0.08 / 0.23, 999 permutations, n = 21). However, the degree of gene reuse was significantly higher for comparisons within a genus (*Arabidopsis*) than between genera (D = 15.37, df = 1, p < 0.001, GLM with binomial errors) while such a trend was absent for parallel function candidates (D = 0.38, df = 1, p = 0.54), suggesting that there are limits to gene reuse at above genus-level divergences. Taken together, these results suggest that there are likely similar functions associated with alpine adaptation among different lineages, species and even genera from distinct tribes of Brassicaceae. However, the probability of reusing the same genes within these functions decreases with increasing divergence among the lineages, thus reducing the chance to identify parallel genome evolution.

### Probability of allele reuse underlies the divergence-dependency of gene reuse

Repeated evolution of the same gene in different lineages could either reflect repeated recruitment of the same allele from a shared pool of variants (‘allele reuse’) or adaptation via alleles representing independent mutations in each lineage (*‘de-novo* origin’) (46). To ask if varying prevalence of these two evolutionary processes could explain the observed divergence-dependency of gene reuse, we quantified the contribution of allele reuse vs. *de-novo* origin to the gene reuse in each pair of *A. arenosa* and *A. halleri* lineages and tested whether it scales with divergence.

For each of the 151 parallel gene candidates, we inferred the most likely source of its candidate variant(s) by using a designated likelihood-based modeling approach that investigates patterns of shared hitchhiking from allele frequency covariance at positions surrounding the selected site (DMC method (46)). We contrasted three models of gene reuse, involving (1) selected allele acquired via gene flow, (2) sourced from ancestral standing variation (both 1 and 2 representing allele reuse) and (3) *de-novo* origin of the selected allele. In line with our expectations, the degree of allele reuse decreased with divergence (D = 34.28, df = 16, p < 0.001, GLM with binomial errors; Fig. 4a). In contrast, the proportion of variants sampled from standing variation remained relatively high even at the deepest interspecific comparison (43%; Fig. 4a, Fig. S11). The absolute number of *de-novo* originated variants was low across all divergence levels investigated (Dataset S8). This corresponds to predictions about a substantial amount of shared variation between related species with high genetic diversity (34) and frequent adaptive trans-specific polymorphism in *Arabidopsis* (10, 50–52). Absence of interspecific parallelism sourced from gene flow was in line with the lack of genome-wide signal of recent migration between *A. arenosa* and *A. halleri* inferred by coalescent simulations (Fig. S12).

**Figure 4:**
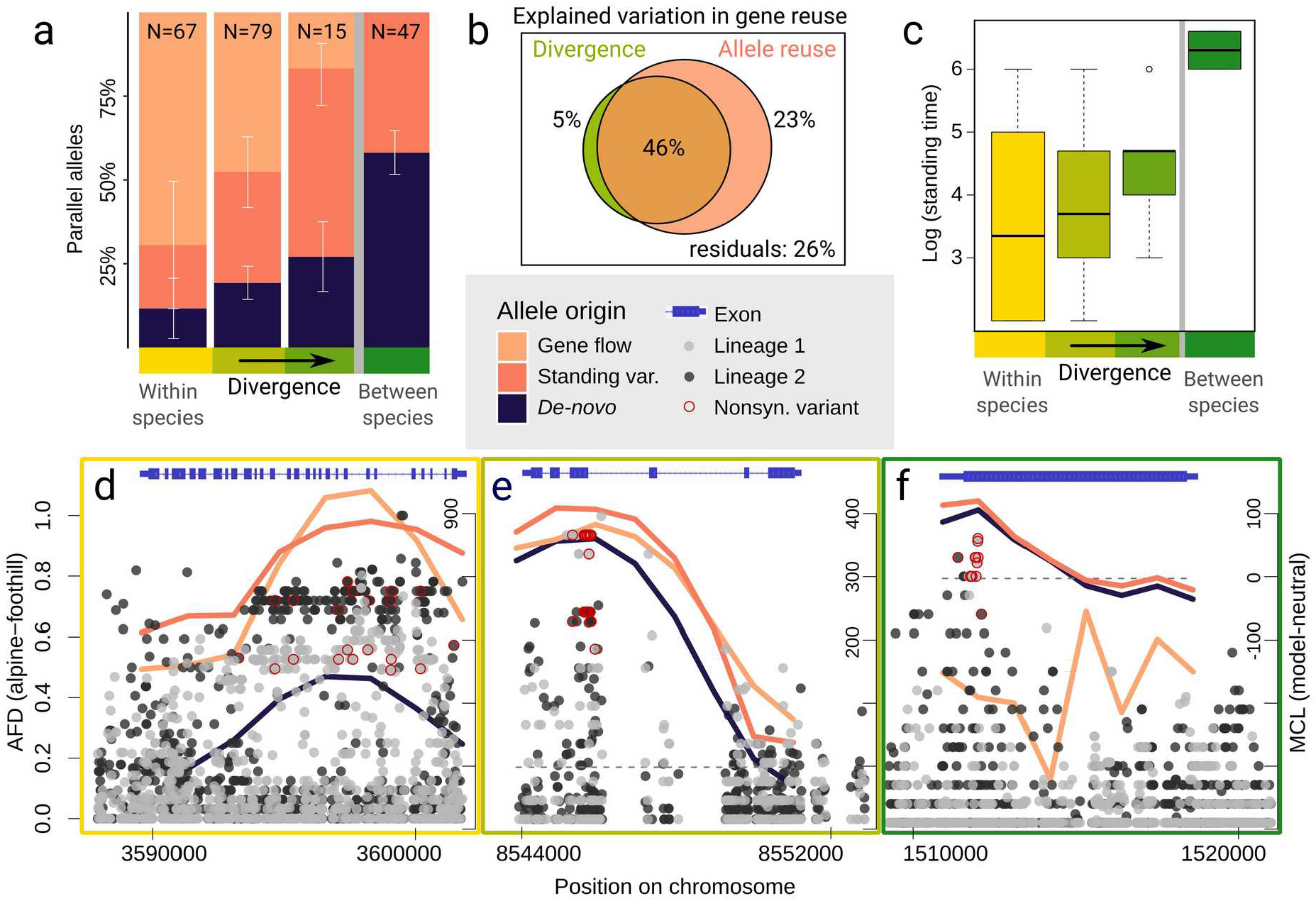
Decreasing probability of allele reuse with increasing divergence in *Arabidopsis arenosa* and *A. halleri*. (a) Proportion of parallel candidate gene variants shared via gene flow between alpine populations from different lineages or recruited from ancestral standing variation (together describing the probability of allele reuse) and originated by independent *de-novo* mutations within the same gene. Percentages represent mean proportions for lineages of a particular divergence category (color ramp; total number of parallel gene candidates is given within each plot). (b) Explained variation in gene reuse between lineages partitioned by divergence (green circle), allele reuse (orange circle) and shared components (overlaps between them). (c) Maximum composite log-likelihood estimate (MCLE) of median time (generations) for which the allele was standing in the populations prior to the onset of selection. (d-f) Examples of SNP variation and MCL estimation of the evolutionary scenario describing the origin of parallel candidate allele. Two lineages in light and dark gray are compared in each plot. (d) Parallel selection on variation shared via gene flow on gene ALA3, affecting vegetative growth and acclimation to temperature stresses (53). (e) Parallel recruitment of shared ancestral standing variation at gene AL730950, encoding heat shock protein. (f) Parallel selection on independent *de-novo* mutations at gene PKS1, regulating phytochrome B signaling (54); here, *de-novo* origin was prioritized over standing variation model based on very high MCLE of standing time (see Methods). Note that each sweep includes multiple highly differentiated nonsynonymous SNPs; in c and d at the same positions in both population pairs, in line with reuse of the same allele. Dotplot (left y-axis): allele frequency difference (AFD) between foothill and alpine population from each of the two lineages (range 0 – 1 in all plots), lines (right y-axis): maximum composite log-likelihood (MCL) difference from a neutral model assuming no parallel selection (all values above dotted grey line show the difference, higher values indicate higher support for the non-neutral model).

Importantly, allele reuse covered a dominant fraction of the variation in gene reuse that was explained by divergence (Fig. 4b) suggesting allele reuse is the major factor contributing to the observed divergence-dependency of gene reuse. We also observed a strong correlation between divergence and the maximum composite-likelihood estimate of the amount of time the allele was standing in the populations between their divergence and the onset of selection (Pearson’s r = 0.83, p < 0.0001, Fig. 4c). This suggests that the onset of selection pressure (assuming a similar selection strength) likely happened at a similar time point in the past. Altogether, the parallel gene candidates (Figure 4d–f) in the two *Arabidopsis* species likely experienced selection at comparable time scales in all lineages, but the degree of reuse of the same alleles decreased with rising divergence between parallel lineages, which explained most of the divergence-dependency of gene reuse.

## Discussion

By analyzing genome-wide variation over twelve instances of alpine adaptation across Brassicaceae, we found that the degree of gene reuse decreased with increasing divergence between compared lineages. This relationship was largely explained by the decreasing role of allele reuse in a subset of seven thoroughly investigated pairs of *Arabidopsis* lineages. These findings provide empirical support for earlier predictions on genetic parallelism (14, 27), and present a general mechanism that may help explain the tremendous variability in the extent of parallel genome evolution that was recorded across different case studies (1, 13). The decreasing role of allele reuse with divergence well-reflects theoretical and empirical findings that the evolutionary potential of a population depends on the availability of preexisting (standing or introgressed) genetic variation (55–57) and that the extent of ancestral polymorphism and gene flow decreases with increasing differentiation between gradually diverging lineages (34, 58). In contrast, the overall low contribution of *de-novo* originated parallel alleles and generally large and variable outcrossing *Arabidopsis* populations suggest a minor role of mutation limitation, at least within our genomic *Arabidopsis* dataset. In general, our study demonstrates the importance of a quantitative understanding of divergence for the assessment of evolutionary predictability (59) and brings support to the emerging view of the ubiquitous influence of divergence scale on different evolutionary and ecological mechanisms (60).

There are potentially additional, non-exclusive explanations for the observed divergence-dependency of gene reuse, although presumably of much lower impact given the large explanatory power of allele reuse in our system. First, theory predicts that the degree of conservation of gene networks, their functions and developmental constraints decrease with increasing divergence (14, 27). Diversification of gene networks, however, typically increases at higher divergence scales than addressed here (millions of years of independent evolution (27)) and affects parallelism caused by independent *de-novo* mutations (18). We also did not find any evidence that our gene reuse candidates were under weaker selective constraint than other genic loci genome-wide. Nevertheless, we cannot exclude that changes in constraint contribute to the decreasing probability of gene reuse across Brassicaceae, as was also reported in (61). Second, protein evolution studies reported patterns of diminishing amino acid convergence over time due to the decreasing probability of hemiplasy, i.e., the gene tree discordance caused by incomplete lineage sorting and introgression (32, 33, 62). As such a pattern reflects neutral processes and is expected to decrease with time, it can confound the assessment of the level of adaptive convergence (32, 33). However, we accounted for this bias in our sampling and analysis design by considering only genes identified as selection candidates in separate divergence scans that contrasted derived alpine populations by their control foothill counterparts. Third, as genetic divergence often corresponds to the spatial arrangement of lineages (63), external challenges posed by the alpine environment at remote locations may differ. Such risk is, however, mitigated at least in our *Arabidopsis* dataset, as the genomically investigated alpine populations share very similar niches (44).

In contrast, no relationship between the probability of gene reuse and divergence was shown in experimental evolution of different populations of yeast (64), raising a question about the generality of our findings. Our study addresses a complex selective agent (a multi-hazard alpine environment (47)) in order to provide insights into an ecologically realistic scenario relevant for adaptation in natural environments. Results might differ in systems with a high degree of self-fertilization or recent bottlenecks, as these might decrease the probability of gene reuse even among closely related lineages by reducing the pool of shared standing variation (65, 66). Although this is not the case in our *Arabidopsis* outcrossers, encompassing highly variable and demographically stable populations, drift might have contributed to the low number of overlaps in comparisons involving the less-variable selfer *Arabidopsis thaliana (42)* in our meta-analysis (Fig. 3e). However, considering the supporting evidence from the literature (Fig. 1a, Fig. S1), and keeping the aforementioned restrictions in mind, we predict that our novel findings are widely applicable. In summary, our study demonstrates divergence-dependency of parallel genome evolution between different populations, species and genera and identifies allele reuse as the underlying mechanism. This indicates that the availability of genomic variation preexisting in the species may be essential for (repeated) local adaptation and consequently also for the predictability of evolution, a topic critical for pest and disease control as well as for evolutionary theory.

## Materials and methods

### Sampling

*Arabidopsis arenosa* and *A. halleri* are biannual to perennial outcrossers closely related to the model *Arabidopsis thaliana*. Both species occur primarily in low to mid elevations (to ~ 1000 m a.s.l.) across Central and Eastern Europe but scattered occurrences of morphologically distinct populations have been recorded from treeless alpine zones (> 1600 m) in several distinct mountain regions in Central-Eastern Europe (43, 67) that were exhaustively sampled by us (Fig. 1, details provided in SI Appendix, Methods).

Here, we sampled and re-sequenced foothill (growing in elevations 460-980 m a.s.l.) as well as adjacent alpine (1625-2270 m a.s.l.) populations from all known foothill-alpine contrasts. In total, we sequenced genomes of 111 individuals of both species and complemented them with 63 published whole genome sequences of *A. arenosa* (68) totaling 174 individuals and 22 populations (Table S1). Ploidy of each sequenced individual was checked using flow cytometry following (69).

### Sequencing, raw data processing, variant calling and filtration

Samples were sequenced on Illumina HiSeq X-ten, mapped to reference genome *Arabidopsis lyrate* (71) and processed following (68). Details are provided in SI Appendix, Methods.

### Population genetic structure

We calculated genome-wide 4d within-(nucleotide diversity (π) and Tajima’s D (70)) and between-(F_ST_ (71)) population metrics using python3 ScanTools_ProtEvol pipeline (github.com/mbohutinska/ScanTools_ProtEvol). ScanTools_ProtEvol is a customized version of ScanTools, a toolset specifically designed to analyze diversity and differentiation of diploid and autotetraploid populations using SNP data (68). To overcome biases caused by unequal population sizes and to preserve the most sites with no missing data, we randomly subsampled genotypes at each position to six individuals per population.

We quantified divergence between pairs of lineages as average pairwise 4d-F_ST_ between the foothill populations as they likely represent the ancestral state within a given lineage. To control for potential effects of linked selection on our divergence estimates, we also extracted all putatively neutral sites that are unlinked from the selected sites (i.e., sites > 5 kb outside genic and conserved regions and sites > 1 Mb away from the centromere). We found out that the genome-wide pairwise inter-population F_ST_ calculated using such sites strongly correlated with 4d-F_ST_ (Pearson’s r = 0.93, p-value < 0.001) and therefore we used only 4d-F_ST_ in further analyses of population structure.

Next, we inferred relationships between populations using allele frequency covariance graphs implemented in TreeMix v. 1.13 (72). We ran TreeMix allowing a range of migration events; and presented two and one additional migration edges for *A. arenosa* and *A. halleri*, as they represented points of log-likelihood saturation (Fig. S4). To obtain confidence in the reconstructed topology, we bootstrapped the scenario with zero events (the tree topology had not changed when considering the migration events) choosing a bootstrap block size of 1000 bp, equivalent to the window size in our selection scan, and 100 replicates. Finally, we displayed genetic relatedness among individuals using principal component analysis (PCA) as implemented in adegenet (73).

We further investigated particular hypotheses regarding the demographic history of our system using coalescent simulations implemented in fastsimcoal2 (74). We calculated joint allele frequency spectra (AFS) of selected sets of populations from genome-wide 4d-SNPs and compared their fit to the AFS simulated under different demographic scenarios. We used wide range of initial parameters (effective population size, divergence times, migration rates, see attached *est* file, Dataset S10). Details on demography inference using fastsimcial2 and coalescent rates in Relate are provided in SI Appendix, Methods.

### Genome-wide scans for directional selection

To infer SNP candidates, we used a combination of two different divergence scan approaches, both of which are based on population allele frequencies and allow analysis of diploid and autopolyploid populations.

First, we calculated pairwise window-based F_ST_ between foothill and alpine population pairs within each lineage, and used minimum sum of ranks to find the candidates. For each population pair, we calculated F_ST_ (75) for 1 kb windows along the genome. Based on the average genome-wide decay of genotypic correlations, (150-800 bp, Fig. S13, SI Appendix, Methods), we designed windows for the selection scans to be 1 kb, i.e., at least 200 bp larger than the estimated average LD. All calculations were performed using ScanTools_ProtEvol, and custom R scripts (github.com/mbohutinska/ProtEvol/selScans.R). Our F_ST_-based detection of outlier windows was not largely biased towards regions with low recombination rate (as estimated based on the available *A. lyrata* recombination map (39) and also from our diploid population genomic data; Fig. S8, S9). This corresponds well with outcrossing and high nucleotide diversity that aids divergence outlier detection in our species (76).

Whenever two foothill and two alpine populations were available within one lineage (i.e., aFG, aNT, aVT and aZT populations of *A. arenosa*), we designed the selection scan to account for changes which were not consistent between the foothill and alpine populations (i.e., rather reflected local changes within one environment). Details are provided in SI Appendix, Methods. Finally, we identified SNPs which were 5% outliers for foothill-alpine allele frequency differences in the above identified outlier windows and considered them SNP candidates of selection associated with the elevational difference in the lineage.

Second, we used a Bayesian model-based approach to detect significantly differentiated SNPs within each lineage, while accounting for local population structure as implemented in BayPass (details in the SI Appendix, Methods) (77).

Finally, we overlapped SNP-candidate lists from F_ST_ and BayPass analysis within each lineage and considered only SNPs which were outliers in both methods as directional selection candidates. We annotated each SNP candidate and assigned it to a gene using SnpEff 4.3 (78) following *A. lyrata* version 2 genome annotation (79). We considered all variants in 5’ UTRs, start codons, exons, introns, stop codons and 3’ UTRs as genic variants. We further considered as gene candidates only genes containing more than five SNP candidates to minimize the chance of identifying random allele frequency fluctuation in few sites rather than selective sweeps within a gene.

For both selection scans, we used relatively relaxed 95% quantile threshold as we aimed to reduce the chance of getting false negatives (i.e. undetected loci affected by selection) whose extent would be later magnified in overlaps across multiple lineages. At the same time, we controlled for false positives by accepting only gene candidates fulfilling criteria of the two complementary selection scans. Using a more stringent threshold of 1% did not lead to qualitatively different results in regards to the relationship between parallelism and divergence (see Supplementary Text 4).

### GO enrichment analysis

To infer functions significantly associated with foothill-alpine divergence, we performed gene ontology enrichment of gene candidates in the R package topGO (80), using *A. thaliana* orthologs of *A. lyrata* genes obtained using biomaRt (81). We used the conservative ‘elim’ method, which tests for enrichment of terms from the bottom of the GO hierarchy to the top and discards any genes that are significantly enriched in descendant GO terms while accounting for the total number of genes annotated in the GO term (80). We used ‘biological process’ ontology and accepted only significant GO terms with more than five and less than 500 genes as very broad categories do not inform about the specific functions of selected genes (FDR = 0.05, Fisher’s exact test). Re-analysis with ‘molecular function’ ontology led to qualitatively similar results (Fig. 14).

### Quantifying parallelism

At each level (gene candidates, enriched GO categories), we considered parallel candidates all items that overlapped across at least one pair of lineages. To test for a higher than random number of overlapping items per each set of lineages (pair, triplet, etc.) we used Fisher’s exact test (SuperExactTest (82) package in R). Next, we calculated the probability of gene-level parallelism (i.e. gene reuse) and functional parallelism between two lineages as the number of parallel candidate items divided by the total number of candidate items between them (i.e., the union of candidate lists from both lineages). We note that the identification of parallel candidates between two alpine lineages does not necessarily correspond to adaptation to alpine environments as it could also reflect an adaptation to some other trigger or to foothill conditions. But our sampling and selection scans, including multiple replicates of alpine populations originating from their foothill counterparts, were designed in order to make such an alternative scenario highly unlikely.

### Model-based inference of the probability of allele reuse

For all parallel gene candidates, we identified whether they indeed support the parallel selection model and the most likely source of their potentially adaptive variant(s). We used the newly developed composite likelihood-based method DMC (Distinguishing among Modes of Convergent adaptation) (46) which uses patterns of hitchhiking at sites linked to a selected locus to distinguish among the neutral model and three different models of parallel selection (considering different sources of parallel variation): (i) on the variation introduced via gene flow, (ii) on ancestral standing genetic variation and (iii) on independent *de-novo* mutations in the same gene (at the same or distinct positions). In lineages having four populations sequenced (aVT, aZT, aFG and aNT), we subsampled to one (best-covered) foothill and one alpine population to avoid combining haplotypes from subdivided populations.

We estimated maximum composite log-likelihoods (MCLs) for each selection model and a wide range of the parameters (Table S10). We placed proposed selected sites (one of the parameters) at eight locations at equal distance apart along each gene candidate sequence. We analyzed all variants within 25 kb of the gene (both upstream and downstream) to capture the decay of genetic diversity to neutrality with genetic distance from the selected site. We used Ne = 800 000 inferred from *A. thaliana* genome-wide mutation rate (83) and nucleotide diversity in our sequence data (Table S4) and a recombination rate of 3.7e^−8^ determined from the closely related *A. lyrata (39)*. To determine whether the signal of parallel selection originated from adaptation to the foothill rather than alpine environment, we ran the method assuming that parallel selection acted on (i) two alpine populations or (ii) two foothill populations. For the model of parallelism from gene flow we allowed either of the alpine populations to be the source of admixture.

For each pair of lineages and each gene candidate, we identified the model which best explained our data as the one with the highest positive difference between its MCL and that of the neutral model.

We further simulated data under the neutral model to find out which difference in MCLs between the parallel selection and neutral model is significantly higher than expected under neutrality. For details see SI Appendix, Methods.

The R code to run the DMC method over a set of parallel population pairs and multiple gene candidates is available at github.com/mbohutinska/DMCloop.

### Statistical analysis

As a metric of neutral divergence between the lineages within and between the two sequenced species (*A. arenosa* and *A. halleri*) we used pairwise 4d-F_ST_ values calculated between foothill populations. These values correlated with absolute differentiation (D_XY_, Pearson’s r = 0.89, p < 0.001) and geographic separation within species (rM = 0.86 for *A. arenosa*, p = 0.002, Fig. 1b) and thus reasonably approximate between-lineage divergence.

To test for a significant relationship between the probability of parallelism and divergence at each level, we calculated the correlation between Jaccard’s similarity in the identity of gene/function candidates in each pair of lineages and (i) the 4d-F_ST_ distance matrix (*Arabidopsis* dataset) or (ii) the time of species divergence (Brassicaceae meta-analysis) using the mantel test with 999 replications (ade4 (84) package in R). Then, we tested whether the relative proportion of the two different evolutionary mechanisms of parallel variation (allele reuse vs. *de-novo* origin) relate to divergence using generalized linear models (R package stats (85)) with a binomial distribution of residual variation. We used the 4d-F_ST_ as a predictor variable and counts of the parallel candidate genes assigned to either mechanism as the explanatory variable. Finally, we used multiple regression on distance matrices (R package ecodist (86)) and calculated the fraction of variation in gene reuse that was explained by similarity in allele reuse, divergence and by their shared component using the original matrices of Jaccard’s similarity in gene and allele identity, respectively, following (87).

### Data Availability

Sequence data that support the findings of this study have been deposited in the Sequence Read Archive (SRA; https://www.ncbi.nlm.nih.gov/sra) with the study codes SRP156117 (released) and SRP233571 (will be released upon publication) (Dataset S9).

## Acknowledgments and funding sources

This manuscript greatly benefited from constructive feedback of Graham Coop, Mike Nowak, Antonín Machač, Anja Westram, Pádraic Flood, Kristin Lee, Timothy Sackton, Martin Weiser and Clément Lafon-Placette. The authors further thank Daniel Bohutínský, Frederick Rooks, Jakub Hojka, Eliška Záveská and Peter Schönswetter for help with field collections; Gabriela Šrámková and Lenka Flašková for help with laboratory work and Doubravka Požárová for help with figure editing. This work was supported by the Czech Science Foundation (GACR project 17-20357Y to FK), a student grant of the Charles University Grant Agency (GAUK 284119 to MB) and long-term research development project No. RVO 67985939 of the Czech Academy of Sciences. MF was supported by a grant from the Swedish Research Council (grant #621-2013-4320 to TS). BL was supported by a grant from SciLifeLab to TS. Sequencing was performed by the Norwegian Sequencing Centre, University of Oslo and the SNP&SEQ Technology Platform in Uppsala. The latter facility is part of the National Genomics Infrastructure (NGI) Sweden and Science for Life Laboratory. The SNP&SEQ Platform is also supported by the Swedish Research Council and the Knut and Alice Wallenberg Foundation. Computational resources were provided by the CESNET LM2015042 and the CERIT Scientific Cloud LM2015085, under the program Projects of Large Research, Development, and Innovations Infrastructures.

## Supplementary Methods

### Sampled populations

Diploid *A. arenosa* populations colonized alpine stands only in Vysoké Tatry (aVT) mountains in Slovakia; more widespread autotetraploids (with random pairing among the four homologous chromosomes (1)) colonized alpine stands in Západné Tatry (aZT) mountains in Slovakia, Făgăraş (aFG) in central Romania, Rodna (aRD) in northern Romania (all these regions are part of the Carpathian Mts.) and Niedere Tauern (aNT) in South-Eastern Alps in Austria (2). Exclusively diploid *A. halleri* species colonized high elevations in Făgăraş (hFG) in Romania and in South-Eastern Alps in Austria (hNT) (3).

The alpine populations in all cases occupy similar habitats (alpine screes and rocky outcrops in glacial cirques, Fig. 1c) and are separated by at least a 500 m altitudinal gap from their foothill counterparts, that also corresponds with timberline (2). Alpine forms of both species are morphologically very distinct from foothill ones but similar together, with lower stature, cushion-like growth form, small, waxy and less hairy leaves and large, usually pinkish petals (Fig. 1d). Although the alpine populations resemble each other phenotypically, they are genetically more closely related to the foothill populations from the same region, suggesting parallel colonization in each range combined with phenotypic convergence (2, 3). Moreover, the widespread occurrence of both species in the foothills vs. rarity in alpine habitats and the fact that the alpine zone of European mountains was previously glaciated suggests that the colonization proceeded from low elevations to the alpine environment (2, 3).

### Sequencing, raw data processing, variant calling and filtration

We extracted DNA from plant material stored in silica gel by CTAB protocol (for details see (4)). Each sample was individually barcoded with dual barcodes during library prep using Illumina TruSeq PCR free kit (Dataset S9), ~ 15 samples were pooled and sequenced on one Illumina HiSeq X-ten lane in a pair-end mode (2× 150 bp) in Norwegian Sequencing Centre, University of Oslo and SNP&SEQ platform in Uppsala.

We used trimmomatic-0.36 (5) to remove adaptor sequences and low quality base pairs (< 15 PHRED quality score). Trimmed reads longer than 100bp were mapped to reference genome *Arabidopsis lyrata* (6) in bwa-0.7.15 (7) with default settings. Duplicated reads were identified by picard-2.8.1 (8) and discarded together with reads that showed low mapping quality (< 25). Afterwards we used GATK (v. 3.7) to call and filter reliable variants and invariant sites according to best practices (9). Namely, we used HaplotypeCaller to call variants per individual with respect to its ploidy level. Then we aggregated variants for all samples per species by GenotypeGVCFs.

We selected only biallelic SNPs and removed those that matched the following criteria: Quality by Depth (QD) < 2.0, FisherStrand (FS) > 60.0, RMSMappingQuality (MQ) < 40.0, MappingQualityRankSumTest (MQRS) < −12.5, ReadPosRankSum < −8.0, StrandOddsRatio (SOR) > 3.0. We called invariant sites using the GATK pipeline similarly to how we did with variants, and we removed sites where QUAL was lower than 15. Both variants and invariants were masked for sites with average read depth higher than 2 times the standard deviation as these sites are most likely located in duplicated regions (duplicated in the genome of our target not in the reference) and regions with excessive heterozygosity, indicating likely paralogous regions mapped on top of each other (genes with > 5 sites which showed fixed heterozygosity in > 2 diploid populations; following (10)). In the final vcf we discarded genotypes with read depth lower than 8 and with more than 20% genotypes missing. Altogether, the dataset contained 11390267 and 6713051 SNPs after variant filtration in *A. arenosa* and *A. halleri*, respectively (Table S2), and the average depth of coverage over both datasets was 32ˣ (Dataset S9).

### Coalescent simulations in fastsimcoal2

First, we tested for the presence of a bottleneck associated with alpine colonization by comparing models with and without a bottleneck. For this analysis we focused on the outlier population with the highest positive 4d-Tajima’s D (i.e., indicative of a bottleneck; LAC from aFG region) and constructed joint AFS using its foothill counterparts from the same (aFG) and next closest (aRD) region (Fig. S6). Second, we tested whether *A. halleri* and *A. arenosa* alpine populations growing in the same mountain regions (i.e. aNT-hNT and aFG-hFG) had experienced recent interspecific gene flow since the last glacial maximum (LGM; approx. 10,000 generations ago assuming a generation time of two years), since their current alpine range was de-glaciated. We constructed the joint AFS from population pairs, iterating over all four combinations of sympatric alpine populations and compared models with recent (post-LGM) migration, ancient (pre-LGM) migration, migration in both periods and without migration (Fig. S12; for *tpl* and *est* files see Dataset S10).

For each scenario and joint AFS, we performed 50 independent fastsimcoal runs with default parameters and n = 10,000, N = 100,000, L = 40, M = 0.001 and l = 10. We then extracted the highest likelihood partition for each fastsimcoal run, calculated Akaike information criterion (AIC) and summarized them across the 50 different runs, over the scenarios and different population pairs/trios. The scenario with the lowest mean AIC values was selected.

### Coalescent rates in Relate

We used program Relate (11) to further refine the demographic history of our system by leveraging haplotype data of diploid populations of *A. arenosa* (aVT lineage) and *A. halleri* (hFG and hNT lineages). First, we phased diploid populations (separately for each species) using the program Shapeit v. 2 (12). We used read aware phasing that accounts for phase information present in the filtered sequencing reads. We also used genetic map of *A. lyrata* as input information (13). We took the phased data from Shapeit and oriented derived alleles based on the polarization table (adopted from (10)). Then we used the command RelateFileFormats to generate a distance file that accounts for uncalled sites. We ran the program Relate per individual chromosome (--mode All) to estimate genome wide genealogies with population size parameter -N set to 3.2e^6^ and mutation rate -m set to 4.3e^−8^. We used the output of the previous command (anc and mut) to estimate coalescence rates between and within populations (script EstimatePopulationSize.sh) in nine iterations with generation time set to 2 years. We ran the commands for each scaffold separately and then we did joint estimation based on information from all the scaffolds. Relative pairwise coalescence was calculated as the proportion of cross-coalescence rate between populations and intra-coalescence rate of one of the populations. All of these estimates were analyzed and visualized in the R script plot_population_size_modif.R.

### Strength of genotypic associations

To design optimal window sizes for the F_ST_ scans, we calculated the strength of genotypic associations (a proxy of linkage disequilibrium, LD) over a range of distances between 10 bp and 50 kb for diploid populations of *A. arenosa* and *A. halleri*. We used PLINK (14) version v1.9 and function r2 (plink --vcf data/scaffold_*.vcf.gz --r2 --ld-window-kb 50 --ld-window 10 --ld-window-r2 0.001 --maf 0.05 --out results/scaffold_* --threads 4 --allow-extra-chr) and summarized the output using custom R scripts. We estimated the average LD as the distance at which genotypic correlation became constant.

### F_ST_ scan for directional selection

We designed the F_ST_ selection scan to account for local changes within each ecotype. To do so, we divided the six possible contrasts among the four populations to four positive (alpine population contrasted with foothill) and two negative (alpine-alpine and foothill-foothill) contrasts. We assigned a rank to each window in each positive contrast, based on the value of F_ST_ (windows with the highest F_ST_ were assigned with the lowest rank) and summed up the ranks over the four positive contrasts per window. We identified windows with the lowest sum of ranks (top 5% outliers of minimum rank sum) as candidates for directional selection. To exclude the local changes uninformative about selection between the foothill and alpine environment, we further identified windows which were the top 5% outliers of F_ST_ in either of the negative contrasts and removed them from the candidate list. For three lineages with only one population pair available (aRD, hNT, hFG), we considered only positive contrast to detect 5% minimum rank outlier windows. We did not observe a decrease in numbers of parallel candidates for between-lineage comparisons including aRD in the same divergence category in *A. arenosa*, suggesting that using two, instead of four populations did not bias our detection of directional selection candidates (Fig. 3a).

### BayPass

We extracted all variable sites for all populations within a particular lineage and calculated reference and alternative allele counts at each site in each population. Then we ran a core model of BayPass which estimates a covariance matrix (approximating the neutral structure among the populations) and differentiation (XtX measure) between populations. We used the default settings; 5000 iterations for the burn-in period, 25000 iterations after burn-in, recorded each 25th (i.e. size of thinning was 25) and 20 pilot runs to adjust the parameters of the MCMC proposal distributions. Then we simulated a set of “neutral” allele counts of 1000 alleles based on our estimated covariance matrix (function simulate.baypass in baypass_utils.R script from BayPass) and ran the same model on the simulated data. We estimated 95% quantile of “neutral” XtX statistics calculated from simulated data and used it to identify excess differentiation SNP candidates in our real dataset.

### Model-based inference of the probability of allele reuse

We simulated data under the neutral model in DMC to find out which difference in MCLs between the parallel selection and neutral model is significantly higher than expected under neutrality. To do so, we generated a distribution of differences between selection model MCLs and the neutral MCL by analyzing neutral data sets, simulated with ms (15), that had similar numbers of segregating sites and demographic history as our real data. We considered the MCL difference between a parallel and neutral model significant if it was higher than the maximum of the distribution of the differences from the simulated data (i.e. 21, a conservative estimate, Fig. S15). We further focused on cases of parallel adaptation in alpine populations by interpreting only cases (loci) for which the model of parallel selection in the alpine environment had a higher MCL than in the foothill environment.

Then, for all genes with significant parallel alpine selection (151 in total, 208 cases of parallel selection between a pair of lineages) we inferred the most likely source of the parallel variant. Analytical theory and simulations show that the ability to distinguish between the standing variation model from the *de-novo* mutation and gene flow models based on MCL is limited at specific parameter combinations (16). Specifically, this depends on the parameter in the standing variation model that specifies the amount of time the allele was standing between the divergence of selected populations and the onset of selection. One cannot distinguish the standing variation model from the *de-novo* mutation model when the maximum likelihood (ML) standing time is much longer than the divergence time of parallel lineages. The standing variation and gene flow models are indistinguishable when the ML standing time is much shorter than the divergence time between the populations experiencing selection (16). Therefore, to categorize all genes into one of the three models, we assigned all genes showing highest support for the standing variation model with a ML standing time of more than 100 thousand and 2 millions of generations for within and between species, respectively, to the *de-novo* mutation model of parallelism. These borderline times were selected as three times the mean estimated times of divergence between *A. halleri* and *A. arenosa* (17) and between *A. arenosa* lineages (1, 18). Similarly, all genes showing highest support for the standing variation model with ML standing time less than 1000 and 1 million generations for within and between species, respectively, were assigned to the gene flow model of parallelism. These borderline times were selected as the lowest non-zero standing time parameters at which the models converged.

The borderline times used gave biologically meaningful results as genes with inferred parallel selection from *de-novo* mutations usually included highly differentiated SNPs at different positions in parallel populations while genes under likely parallel selection from standing variation contained highly differentiated SNPs that were identical (Fig. 4d, e). Moreover, using higher or lower cut-off time (Table S15), did not lead to qualitatively different results in regards to the relationship between parallelism and divergence (Supplementary Text 5).

### Integration of candidate lists from *Brassicaceae*

We gathered lists of gene candidates from all available studies focused on alpine adaptation in Brassicaceae (19–24), i.e., adaptation towards treeless high-elevation habitats addressed by studies of whole genome sequence data (Table S8). We unified (merged) the gene candidate lists if multiple altitudinal gradients were screened within a species (except for *A. halleri* from Europe and Japan which were kept as separate units due to extraordinarily high spatial and genetic divergence of these two lineages) (3). We obtained 19 – 1531 *A. thaliana* orthologues and annotated them into functions using gene ontology enrichment in the same way as described earlier.

We note that gene sets identified by the revisited studies may partly differ also due to the different genome properties of diverged species and varying methods used to detect candidate genes in them (for details see Table S8). However, a dramatic rearrangement of gene number and/or gene functions genome-wide is unlikely among species from the same family (25). Moreover, the total numbers of identified candidate genes in individual studies did not decrease with increasing divergence from *Arabidopsis* (e.g. analysis of distantly related *Crucihimalaya himalaica* identified a higher number of candidate genes than analyses of most of the *Arabidopsis* species, Fig. 3e). Thus the pattern of divergence-driven decrease of probability of gene-level parallelism does not reflect the mere limits of candidate detection methods. Still, keeping such potential technical limitations in mind, we did not interpret any specific values obtained from the analysis.

## Supplementary Texts

### Supplementary Text 1: Exclusion of the aVT-aZT lineage pair does not qualitatively affect the results

In the genome-wide analysis of population structure among the lineages (Figs. S3 and S4) we observed grouping of alpine populations from aVT and aZT lineages, suggesting they do not represent entirely independent instances of alpine colonization. However, taking into account the separation of aVT and aZT populations by ploidy (aVT populations are diploid while aZT are tetraploid) and previously detected single origin of the tetraploid *Arabidopsis arenosa*, the scenario of independent alpine colonization followed by later interploidy introgression is likely (1, 26). Thus, we analyzed both regions separately. In order to assess the potential influence of such a decision on our findings, we excluded the aVT-aZT pair here and re-ran the analyses. The results were not different from those presented in the main text (Figs. 3, 4). Re-calculating the results without the aVT-aZT pair, we found that parallelism by gene was less common than by function (mean proportion of parallel items = 0.044 and 0.066, respectively, and this difference was significant, p < 0.001, GLM with binomial errors).

While the proportion of parallels at the gene level was significantly correlated with divergence (negative relationship between Jaccard’s similarity in candidate gene identity among lineages and 4d-F_ST_; Mantel test, rM = −0.68, respectively, p-value < 0.05 in both cases), no relationship was identified for the proportion of parallels by function (rM = 0.11, p-value = 0.6, Mantel test).

Similarly, the results regarding the mechanism of parallelism (allele reuse vs. *de-novo* origin) were not qualitatively affected. With increasing divergence, the proportion of allele reuse decreased (p < 0.001, GLM with binomial errors) and the onset of selection pressure (assuming a similar selection strength) likely happened at a similar time point in the past. This was indicated by a strong correlation between divergence and the maximum composite-likelihood estimate of time for which the allele was standing in the populations prior to the onset of selection (Pearson r = 0.80, p < 0.001, log-transformed time).

### Supplementary Text 2: Parallel gene candidates were not outliers from the genome-wide background

The parallel gene candidates were distributed equally across all chromosomes and they did not cluster in regions with extreme values of recombination rate per gene, as estimated based on the available *A. lyrata* recombination map (19) (Fig. S8). Likewise, the peaks of differentiation (F_ST_) around loci of parallel gene candidates did not overlap with regions of low recombination rate, estimated from our diploid population genomic data (Fig. S9). This is consistent with strict outcrossing and high nucleotide diversity in *A. arenosa* and *A. halleri* (27) both reducing the effect of varying recombination rate on F_ST_-based inference of divergence.

Longer genes represent larger mutational targets and thus on average harbor more polymorphism. Thus, we next checked for differences in distributions of genome-wide and parallel candidate gene lengths. In line with this hypothesis, parallel candidate genes were on average significantly longer than expected given the genome-wide distribution of gene lengths (Fig. S16, Table S11). However, this difference was rather subtle and parallel candidates were not strong outliers for this metric (only a single parallel candidate gene, AL420810, 21 kb, fell within an extreme 0.1% tail of genome-wide distribution of gene lengths).

Genes where mutations have less negative pleiotropic effects (i.e. with weaker selective constraints) could be more likely to become parallel candidates. Thus, we estimated the degree of the constraints in 50-kb windows in foothill populations of the seven lineages using the ratio of nonsynonymus over synonymous diversity (π_N_/π_S_). Elevated value of π_N_/π_S_ suggests that the loci are likely less constrained; thus we compared π_N_/π_S_ of windows overlapping with parallel gene candidates (parallel windows) to windows overlapping with other genes genome-wide (nonparallel windows). We did not detect a difference in π_N_/π_S_ of parallel and nonparallel windows in any of our lineages (p = 0.55 – 0.95, Wilcoxon rank test, n of windows = 1780 – 2741, Table S7, Fig. S7, Table S6) and thus we found no evidence that variability in constraints on gene evolution is affecting the probability of parallel gene reuse in our dataset.

### Supplementary Text 3: Putative alpine adaptations and their functional implications

To characterise likely functions of the set of parallel gene candidates, we first annotated each of them using *A. thaliana* gene function descriptions from the TAIR database and targeted search in the publications associated with the genes in the TAIR, if needed. Out of the 151 parallel gene candidates, we managed to annotate 146, 13 of which were annotated with unknown function, leaving us with 133 informative functional annotations (Dataset S2). Second, we searched for possible interactions among the set of parallel gene candidates, using protein-protein association information from the STRING database (genes identified as associated were further referred to as ‘interacting’). We identified three clusters or interacting genes and six interacting gene pairs (Fig. S10). Third, we identified 22 overrepresented gene functions using GO enrichment analysis (Dataset S3).

Altogether, these analyses suggest that alpine adaptation has a complex genetic basis, including several genes and functions, which is to be expected for a response to a multifactorial environmental stress.

We divide the functions of candidate genes into two main categories.

#### 1. Physiological response to alpine stresses

##### Abiotic stresses

###### Cold (drought) response

In many aspects, plants exhibit common molecular and physiological responses on exposure to cold and drought. Both factors result in water deficit and following turgor stress and membrane damage. Drought and chilling also exacerbates ROS production in plants’ cells, causing oxidative stress. Multiple enriched GO terms relate to these types of stresses, including “stomatal movement”, “cellular response to external stimulus”, “cellular response to environmental stimulus” and “response to wounding”. Related candidate genes involve ALA3, a phospholipid translocase which has been associated with adaptability to temperature stresses, and its third-shell interactor AT3G53830, which was shown to regulate freezing tolerance and cold acclimation through modulating lignin biosynthesis. Further related candidates include LRK10L1.2, which mediates ABA signaling and drought resistance, FAR5, preventing root wound damage and water control by synthesis of suberin and MAP18, expressed in roots and mediating drought and chilling tolerance. Mutants of ACC1, whose product is essential for very long chain fatty acid elongation, are deficient in freezing tolerance after cold acclimation.

**Response to high light stress/reactive oxygen species** is represented by enriched GO terms “response to blue light”, “cellular response to environmental stimulus” and “response to decreased oxygen levels”. It involves PAP1, one of the candidates with the highest number of differentiated SNPs, responsible for anthocyanin biosynthesis under red light and up-regulated by genotoxic stress (gamma ray or UV-B). A pair of interacting genes CH1, involved in response to high light stress by chlorophyll biosynthesis, and AT5G14700, involved in gamma irradiation response through lignin biosynthesis. A pair of interactors AT5G22620 and AT5G65750, reacting to oxidative stress damage. Further, G3Pp3 is reacting to gamma irradiation, AHG11 mutants have higher levels of reactive-oxygen-species-responsive genes and CHR8 is involved in DNA repair in response to gamma radiation.

**Timing of life cycle, mainly flowering and circadian rhythms**, is represented by enriched GO terms “response to blue light”, “regulation of flower development” and “cellular response to external stimulus”. One of our candidate genes is FT, which acts together with CONSTANS as the main regulator of vernalization and flowering time. Our list further includes homolog of CONSTANS COL1, PKS1, which is a negative regulator of phyB signalling and TZP, regulator of circadian rhythms. TIL2 is related to pollen germination and AT1G52060 to seed germination.

**Ion transport** is represented by GO terms “cellular response to nitrogen compound”, “cellular chemical homeostasis” and “inorganic anion transport”. It involves number of candidate genes, among them NRT2;1, GLR2.5 and WR3, involved in nitrate transport, sulphate transporter SULTR1;1, NCL, regulating calcium homeostasis, AT5G65750, involved in zinc and cadmium binding, a pair of interacting genes STOP1 and ABCB27, and PAH2, involved in aluminium sensitivity or ABCG37, VIT and AT5G14700, which are related to iron homeostasis.

##### Biotic stresses

**Response to pathogens and herbivory** is represented by multiple significantly enriched GO terms, including “leaf senescence”, “response to wounding”, “beta-glucan metabolic process”, “glucosinolate metabolic process” and “glycosyl compound metabolic process”. In *A. thaliana*, glucosinolate metabolism products support a broad spectrum of pathogenic infection responses and are associated with protection against herbivores. Numerous candidate genes are involved in glucosinolate metabolism and pathogen defense, including GH9B10, ESP, AT4G10060, AT5G25980, FST1, UGT85A3 and AT5G59530. Wound response is mediated via GLR2.5, involved in wound signaling, WOUND-RESPONSIVE 3 (WR3), which is mainly responsible for nitrate uptake but also to root wounding response, an aromatic aldehyde synthase AAS, a plasma membrane localized ATPase transporter MRP4 and FAR5, which generate the fatty alcohols found in root, seed coat, and wound-induced leaf tissue. Herbivore resistance is further mediated by a chitinase AT2G43570. Response to foreign attacks by pathogens is mediated by a family of wall-associated kinases, represented by WAK2, WAKL4 and WAKL5 and defense signaling pathway gene 5.

#### 2. Morphological changes related to the alpine adaptation

##### Root growth and root hair production

A trio of interacting genes ABCG37, AT1G52060 and MAP18 are all overexpressed in roots, MAP18 is directly linked to root hair elongation. Mutants of ALA3 have short primary roots. Roots are further affected by a pair of interactors AT5G49770, responsible by root hair development and SUS4, affecting root growth under hypoxia. RSH2 is responsible for root epidermis cell differentiation in Arabidopsis and SABRE for cell expansion in the root cortex. NRT2;1 represses lateral root initiation.

**Flower, gamete and seed development** is represented by significantly enriched GO term “regulation of flower development”. SEU mutants have altered floral identity development and reduced the number of seeds per silique. TIL2 was associated with pollen germination, AT5G03560, GLR2.5, BSL2 and STP8 are related to pollen tube growth, AT4G27290 to pollen recognition. FAR5 contributes to the seed coat barrier formation, AT1G77520 and TT9 to secondary metabolism in seeds, AT4G22520 forms seed storage albumin protein, AT1G52060 is associated with seed germination. TOPP6 is dominantly expressed in siliques.

**Plant size reduction** is related to the enriched GO term “developmental growth involved in morphogenesis”. Further, mutants of ALA3, one of the candidates with the highest number of differentiated SNPs, are characterized by slower vegetative growth. Similarly, SEU mutants show reduced plant height and increased lateral branching. The remaining candidate genes and enriched GO terms are related to general roles in metabolism and signaling, which might play a crucial role in the alpine adaptation, but are difficult to interpret.

### Supplementary Text 4: Re-analysis with 1% threshold did not yield qualitatively different results

The choice of significance threshold for outlier detection in selection scans is an arbitrary decision. In order to detect a high number of true selection events in the genome while keeping the false positive rate low, we used a relatively relaxed 5% threshold but considered only signals which were outliers in a combination of two different divergence scan approaches. Here, we show that analysis with 1% threshold yields results comparable to the findings presented in the main text. Specifically, we found that parallelism by gene was less common than by function (mean proportion of parallel items = 0.022 and 0.044, respectively, and this difference was significant, p < 0.001, GLM with binomial errors).

Similarly, the probability of parallelism at the SNP and gene level significantly correlated with divergence (negative relationship between Jaccard’s similarity in candidate gene identity among mountain regions and 4d-FST; Mantel test, rM = −0.60, p-value < 0.05 in both cases). In addition, we found a significant relationship between divergence and the proportion of parallels by function (rM = −0.66, p-value = 0.01, Mantel test). This effect was, however, driven by the low probability of parallelism by function between species, which itself reflects the low number of enriched GO terms (36 – 103) in this analysis employing more stringent threshold; note that the Mantel test compares similarity in enriched GO terms identity among lineages, thus the significance is always based on the same number of observations (pairwise comparisons, not the total number of enriched GO terms).

Due to the low number of parallel genes identified under the 1% threshold (only 18 parallel genes passed the MCL difference criterion in DMC, none of them between species or among distant lineages within species), we could not test for the relationship between the mechanism of parallelism (allele reuse vs. *de-novo* origin) and divergence. However, all of the 18 parallel genes identified among closely related lineages (F_ST_ 0.07 – 0.14) came from reuse of shared alleles, further supporting our finding that with decreasing divergence, the proportion of parallelism due to allele reuse increases.

### Supplementary Text 5: Re-analysis of mechanism of parallelism using higher and lower borderline times

We quantified the relative importance of allele reuse and *de-novo* origin of parallel alleles in each regional pair, using one higher and one lower borderline time to distinguish between standing variation and gene flow or *de-novo* mutation models. We used the borderline time of 1000 / 1e6 generations within / between species (instead of original 100 / 1e5) to distinguish between gene flow and standing variation model and 1e5 / 2e6 and 3e6 generations within / between species (instead of 1e6 / 2.5e6) to distinguish between standing variation and *de-novo* origin model (Table S10). Using such datasets, we obtained qualitatively identical results compared to the findings in the main text in terms of dependence between the probability of allele reuse and divergence (Fig. 4a). Specifically, with increasing divergence, the probability of allele reuse decreased, no matter which borderline time we used (p < 0.001 in all three cases, GLM with binomial errors). The only actual difference was a 13 % higher support for the gene flow model at the lowest divergences, when applying a higher borderline time (1000 instead of 100 generations) to differentiate gene flow from standing variation models. However, such a change is not substantial for our conclusions as we evaluated both types of allele reuse (gene flow and standing variation) together.

## Supplementary Figures

**Figure S1:**
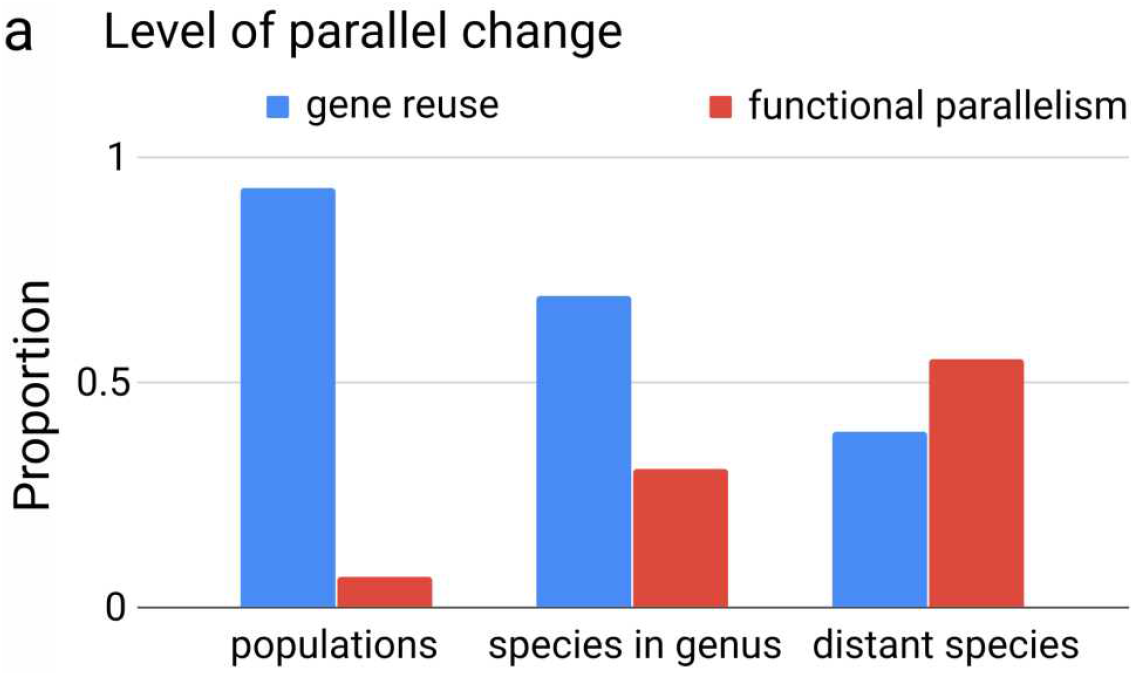
Frequency of gene reuse and functional parallelism inferred from a review of individual case studies categorized based on divergence between parallel lineages. a: Summary over 29 reports show that gene reuse decreases with divergence with a relative increase in parallelism by function. If studies reported a gene reuse leading to functional parallelism, we only classified them as gene reuse. See Dataset S1 for details on the studies reviewed.

**Figure S2:**
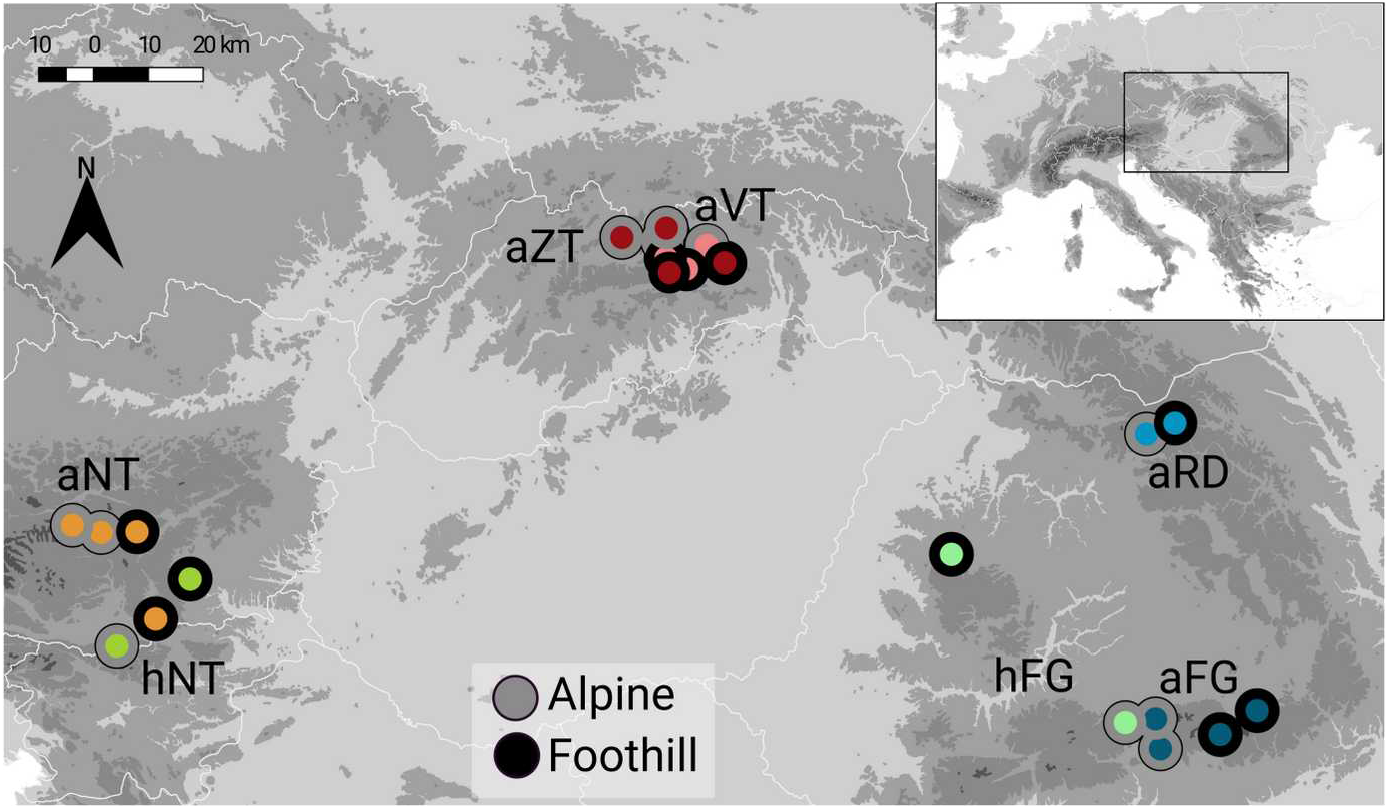
Geographical location of sampled *Arabidopsis arenosa* and *A. halleri* populations. Lineages (*A. arenosa*: aNT, aVT, aZT, aRD and aFG; *A. halleri*: hNT, hFG) are colored by different colors, alpine populations are framed in grey, foothill in black.

**Figure S3:**
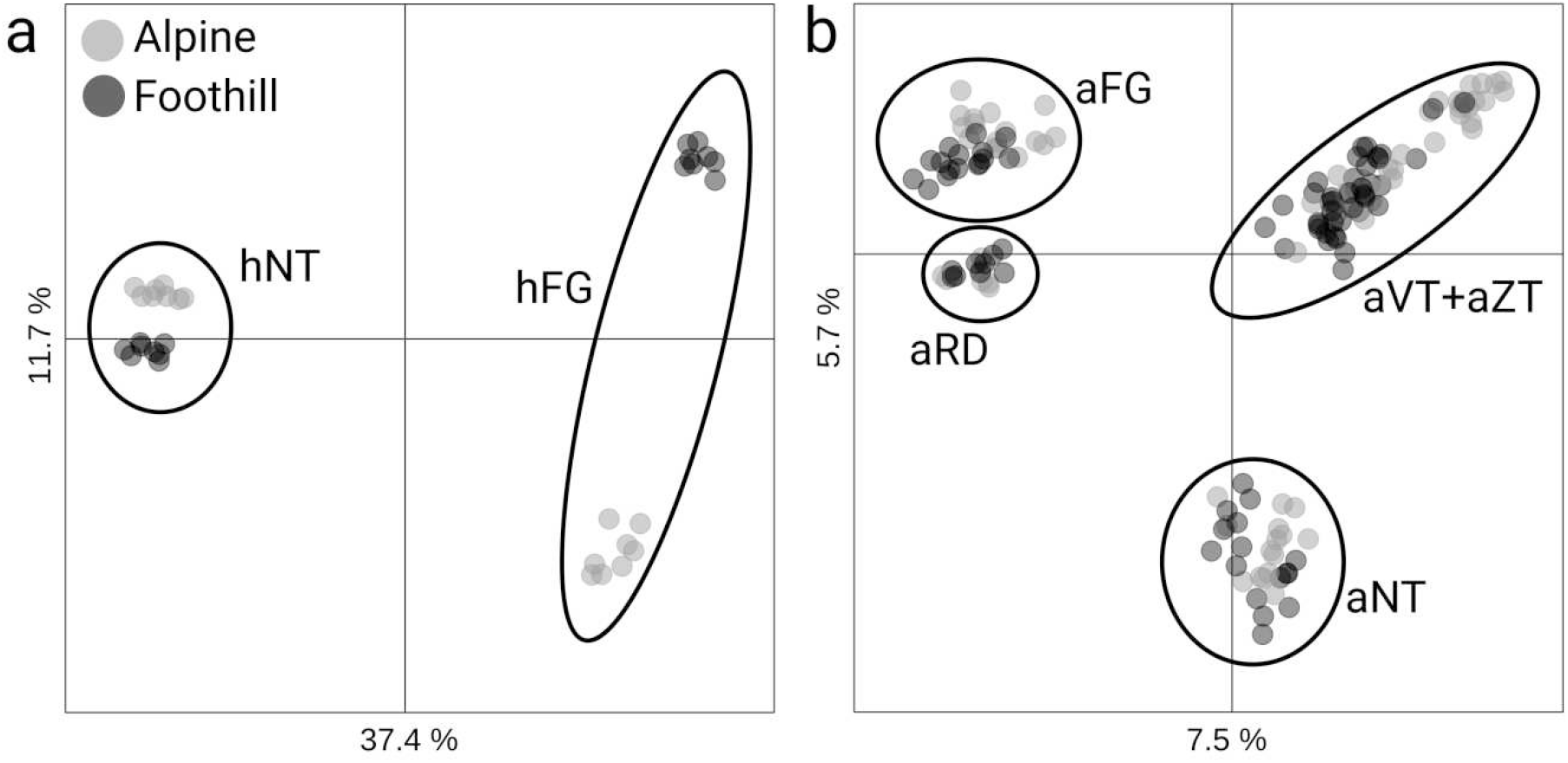
Regional (enclosed with bold lines) rather than elevation-based clustering of alpine (grey) and foothill (black) populations demonstrate repeated origin of parallel alpine ecotype of *Arabidopsis halleri* (a) and *A. arenosa* (b). Shown are first two axes of principal component analysis and percentages of explained variability.

**Figure S4:**
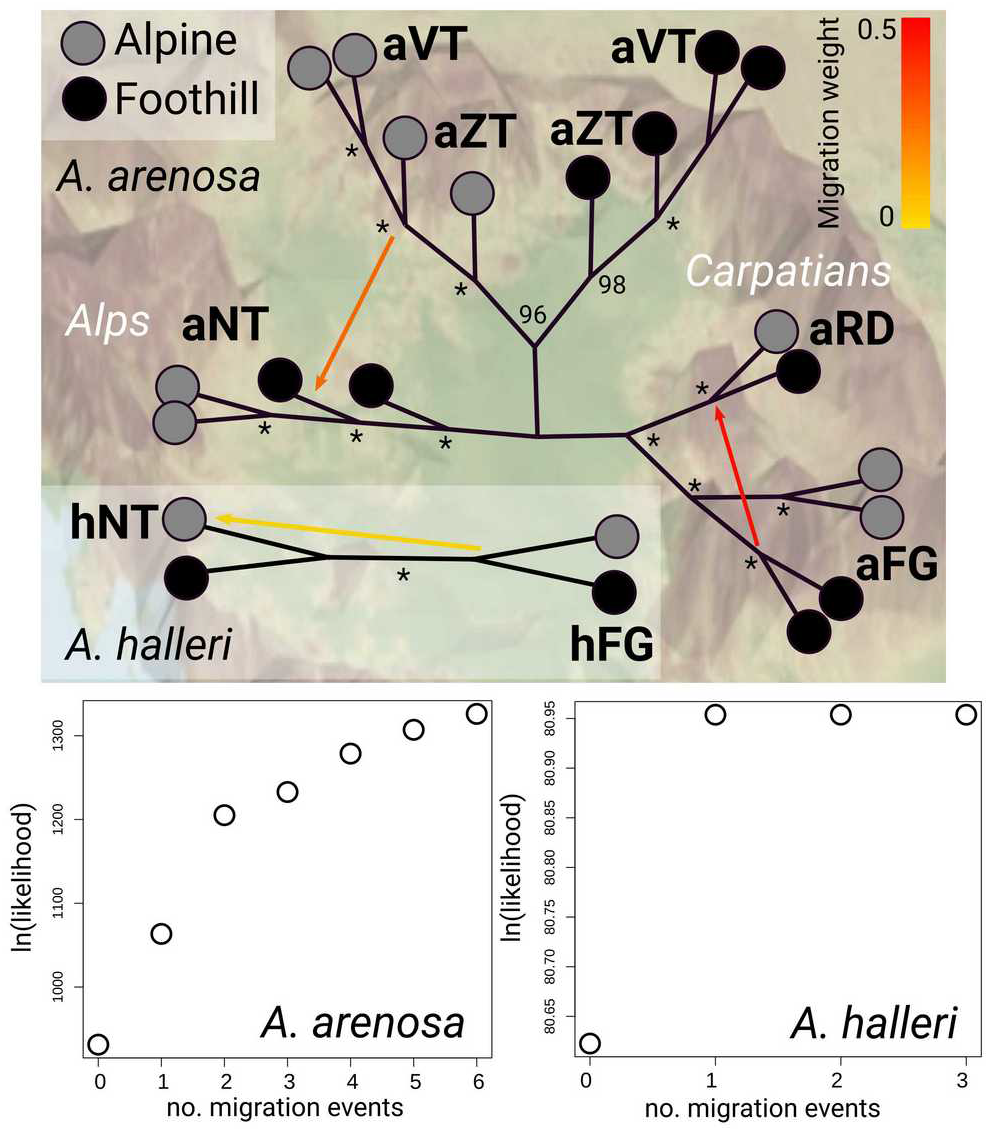
Phylogenetic relationships and migration events between alpine (gray) and foothill (black) populations of *Arabidopsis arenosa* and *A. halleri* inferred by TreeMix. The topology is superimposed onto the geographical distribution of the populations; the asterisks show branch bootstrap support of 100. Bottom part summarizes likelihood at different numbers of allowed migration events; the saturation starts after two and one additional migration edges for *A. arenosa* and *A. halleri*, respectively.

**Figure S5:**
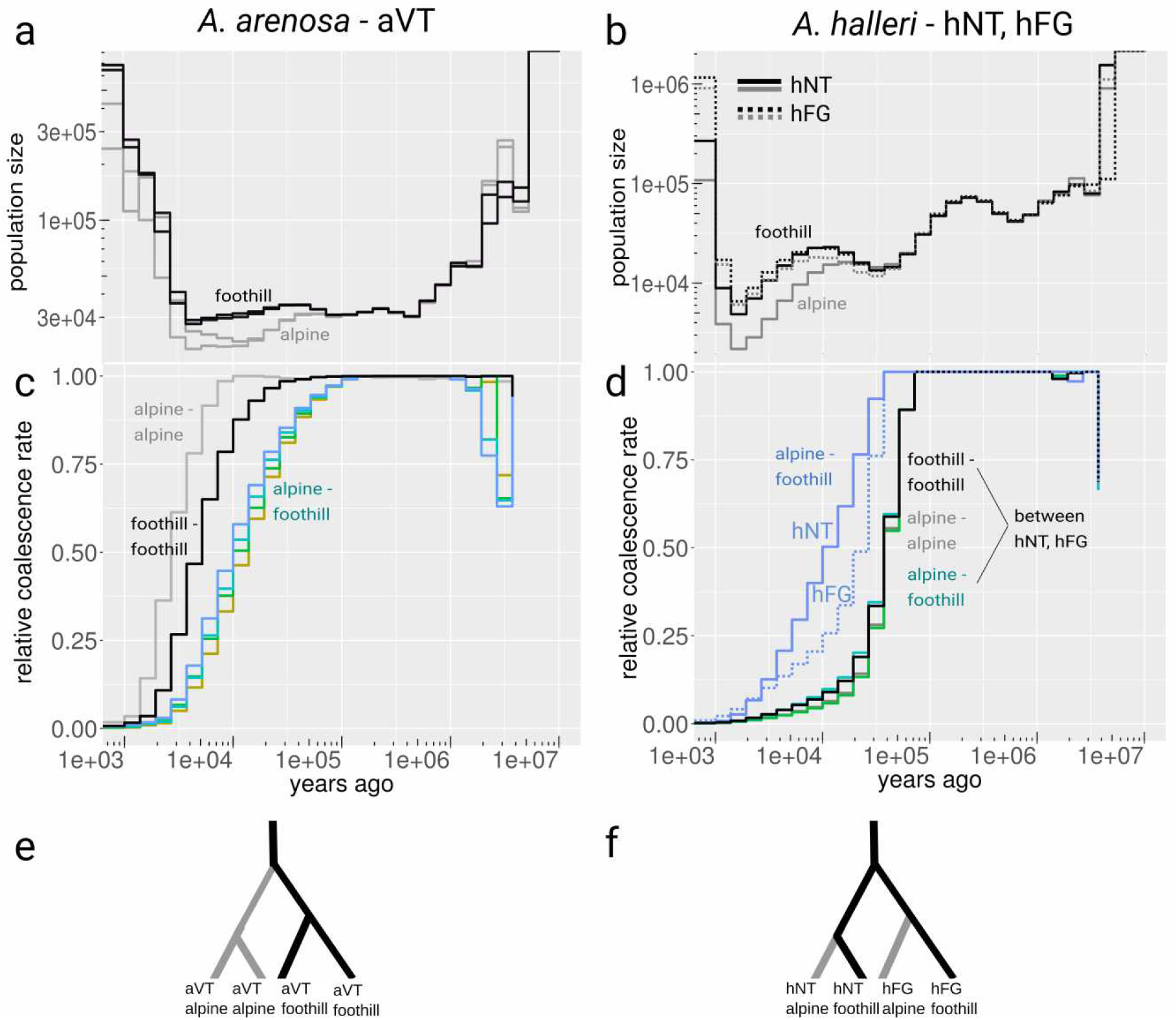
Population sizes and split times in diploid populations of *Arabidopsis arenosa* (aVT lineage, 4 populations) and *A. halleri* (hNT and hFG lineages, 2 populations each) estimated in Relate. a, b: Population-specific estimates of alpine (grey) and foothill (black) population sizes. c, d: Cross-population coalescence rates estimations using genome-wide genealogies between alpine and foothill populations (colored lines), foothill-foothill populations (black lines) and alpine-alpine populations (grey lines). e, f: Interpretation of separation histories between populations based on a, b, c and d. Note that separation histories estimated in Relate are in agreement with population structure analyses based on genome-wide nearly-neutral four-fold degenerate SNPs (Figs. S3 and S4), suggesting parallel alpine colonization of each mountain region by a distinct (foothill) genetic lineage in *A. halleri* (i.e. the only species with multiple regions occupied by diploids). One generation corresponds to two years (assuming biennial life cycle).

**Figure S6:**
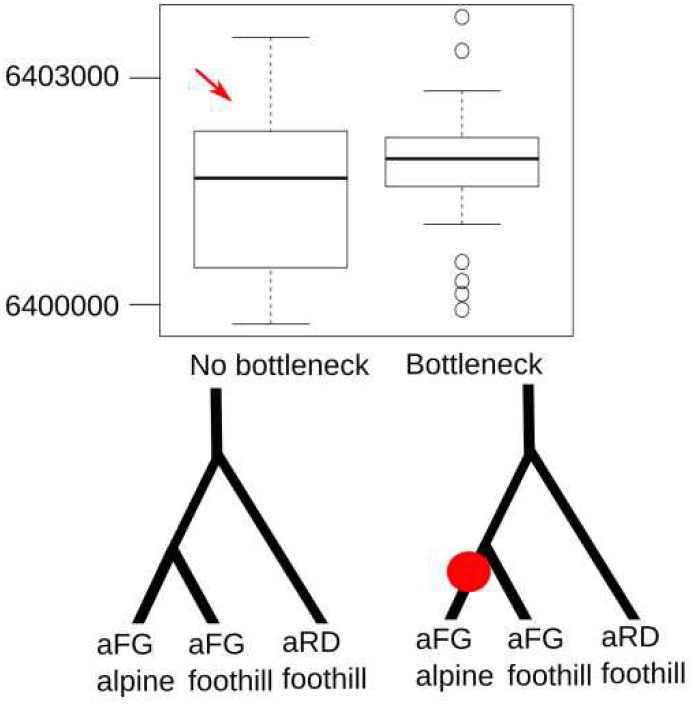
Scenario assuming bottleneck during alpine colonization does not improve model fit for a candidate population. Comparison of Akaike information criteria (AIC) across two scenarios of split of foothill and alpine population from the same lineage without bottleneck and followed by bottleneck. We used the alpine population LAC of aFG which showed the outlying highest Tajima’s D value (0.66, Table S4), i.e., the most probable candidate for a bottleneck. Each scenario was simulated by 50 independent runs of fastsimcoal2, the corresponding distribution of the AIC values over these 50 runs is summarized by the boxplots. Red arrow highlights more likely scenario without bottleneck with significantly lower median AIC (p = 0.03, Tukey HSD test). The simulated topologies and bottleneck (red dot) are depicted below the corresponding plots.

**Figure S7:**
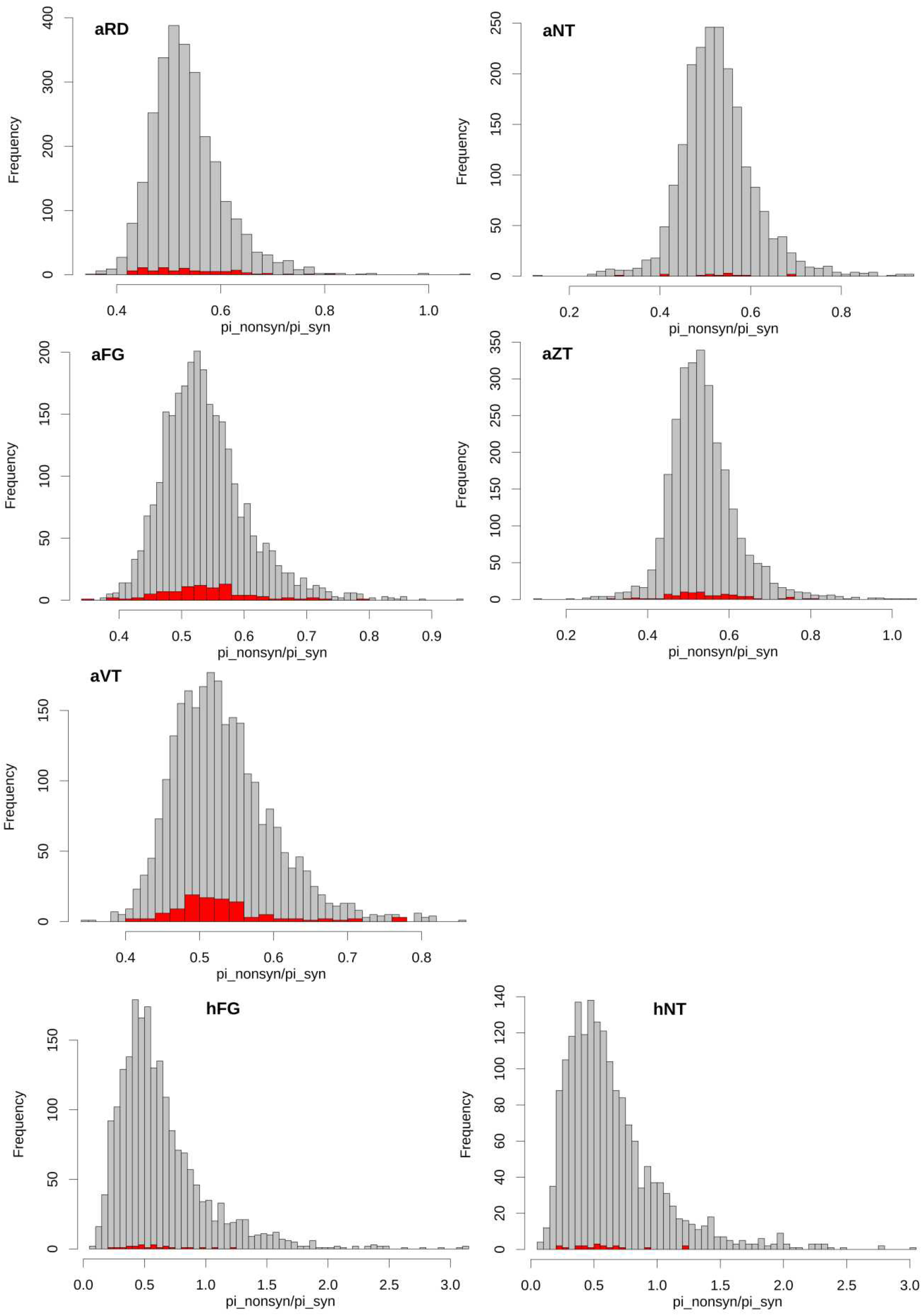
Distribution of π_N_/π_S_ (used as a proxy for selective constraints) in 50-kb windows overlapping with parallel gene candidates (red) and with any other gene in the genome (grey). We found no evidence that windows spanning our parallel candidate genes were subjected to stronger or weaker constraints compared to the rest of the genome (Table S6).

**Figure S8:**
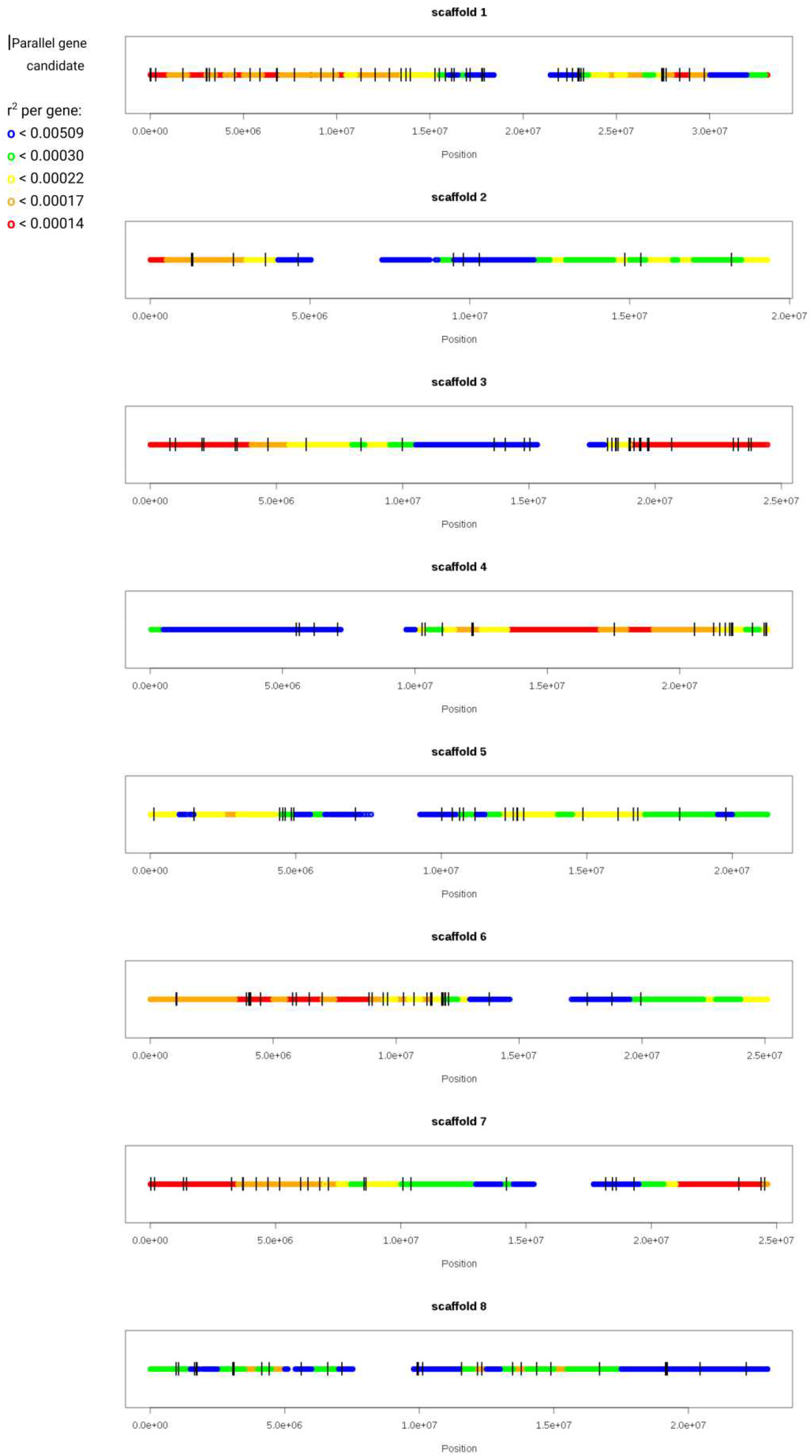
Location of all parallel gene candidates (black vertical lines) on *A. lyrata* reference chromosomes colored by bins of distinct recombination rate per gene. The figure indicates that parallel gene candidates do not cluster in regions with extreme values of recombination rate per gene (blue/green), as estimated based on the available *A. lyrata* recombination map (19).

**Figure S9:**
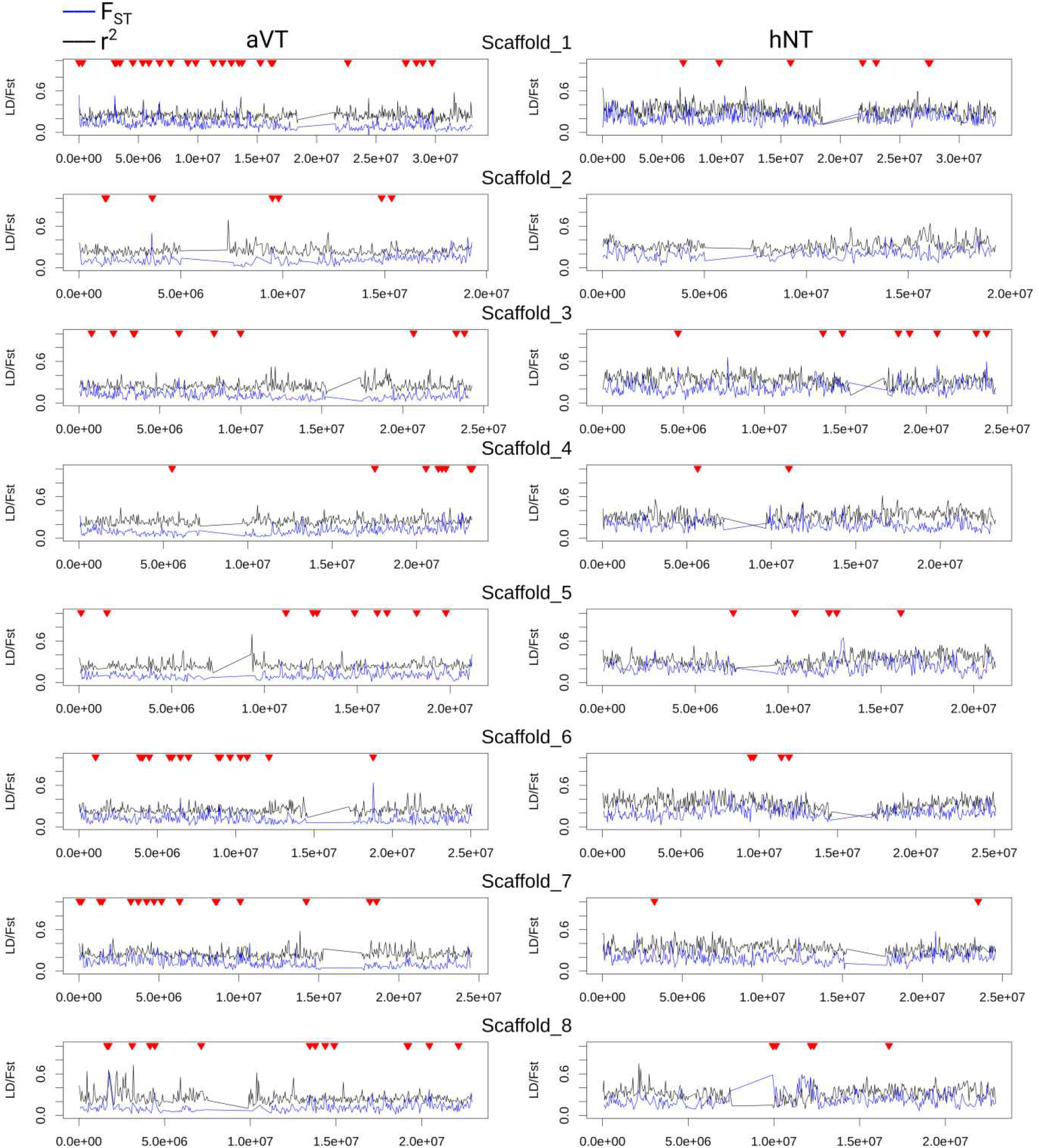
F_ST_ (blue lines) and r^2^ (black lines; proxy for LD) plotted in 50-kb windows along chromosomes. Genetic differentiation (F_ST_) around parallel gene candidates (red triangles) do not correspond to regions of low recombination rate, approximated by regions of high genotypic correlations (r^2^) estimated from our diploid population genomic data from *Arabidopsis arenosa* (aVT, left) and *A. halleri* (aNT, right).

**Figure S10:**
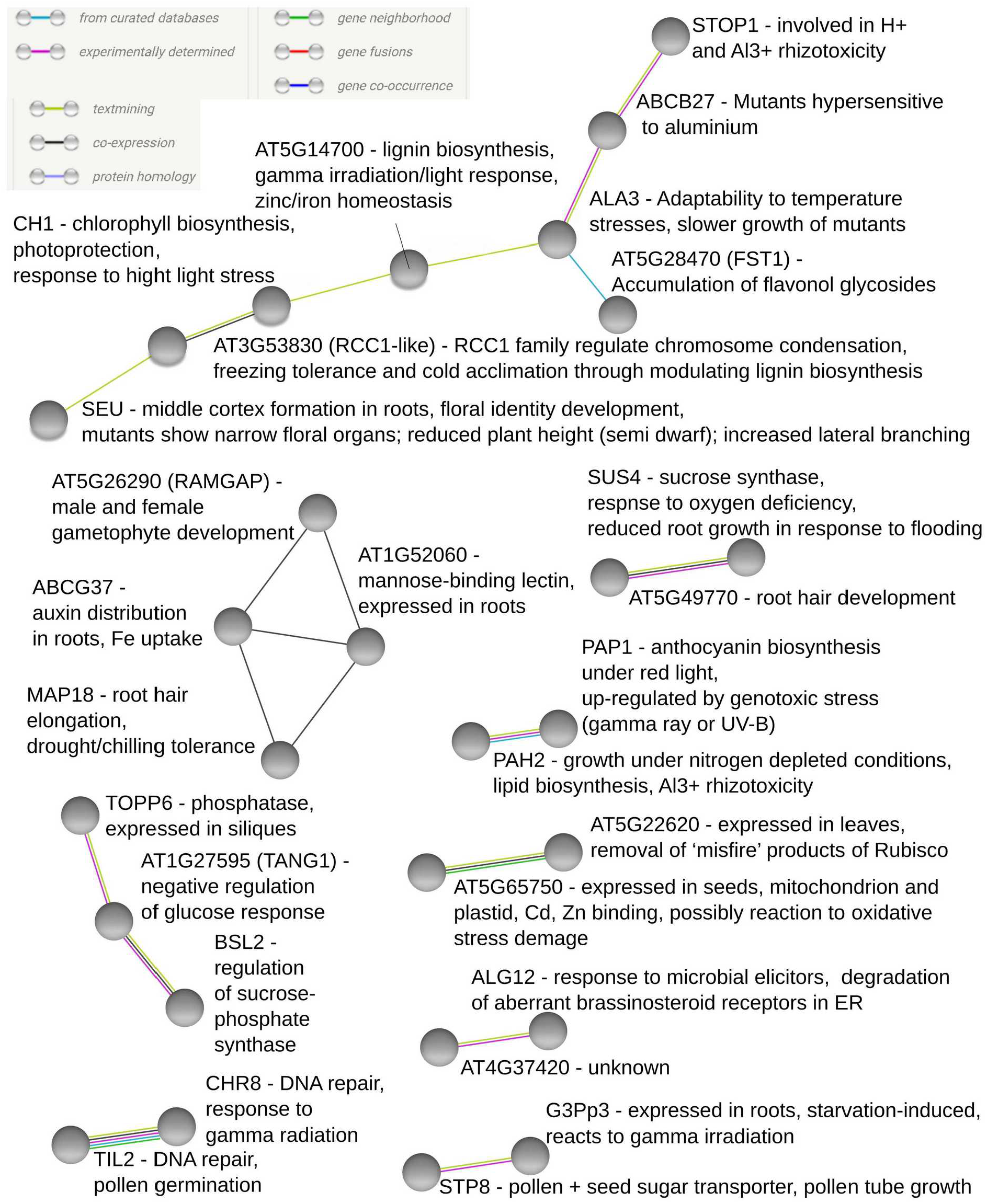
Protein-protein interactions predicted among our set of parallel gene candidates using the STRING and their functional annotations based on the TAIR database and associated literature. We used only medium confidence interactions and higher, colors of edges represent type of evidence supporting each interaction.

**Figure S11:**
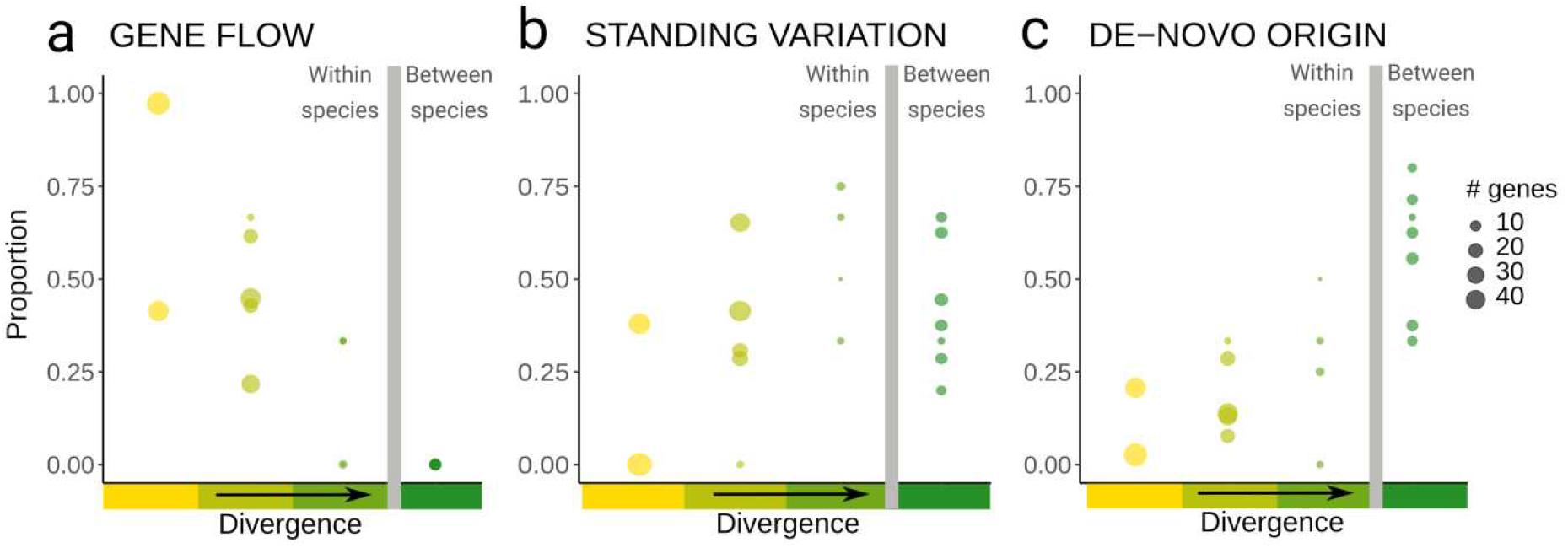
Decreasing probability of allele reuse with increasing divergence among the population pairs. a: alleles reused via gene flow between alpine populations from different regions, b: allele reuse from ancestral standing variation, c: parallel variation from independent *de-novo* mutations targeting the same gene. Proportions are calculated across candidate genes selected in parallel between pairs of populations from different mountain ranges (size of the dot corresponds to the number of genes).

**Figure S12:**
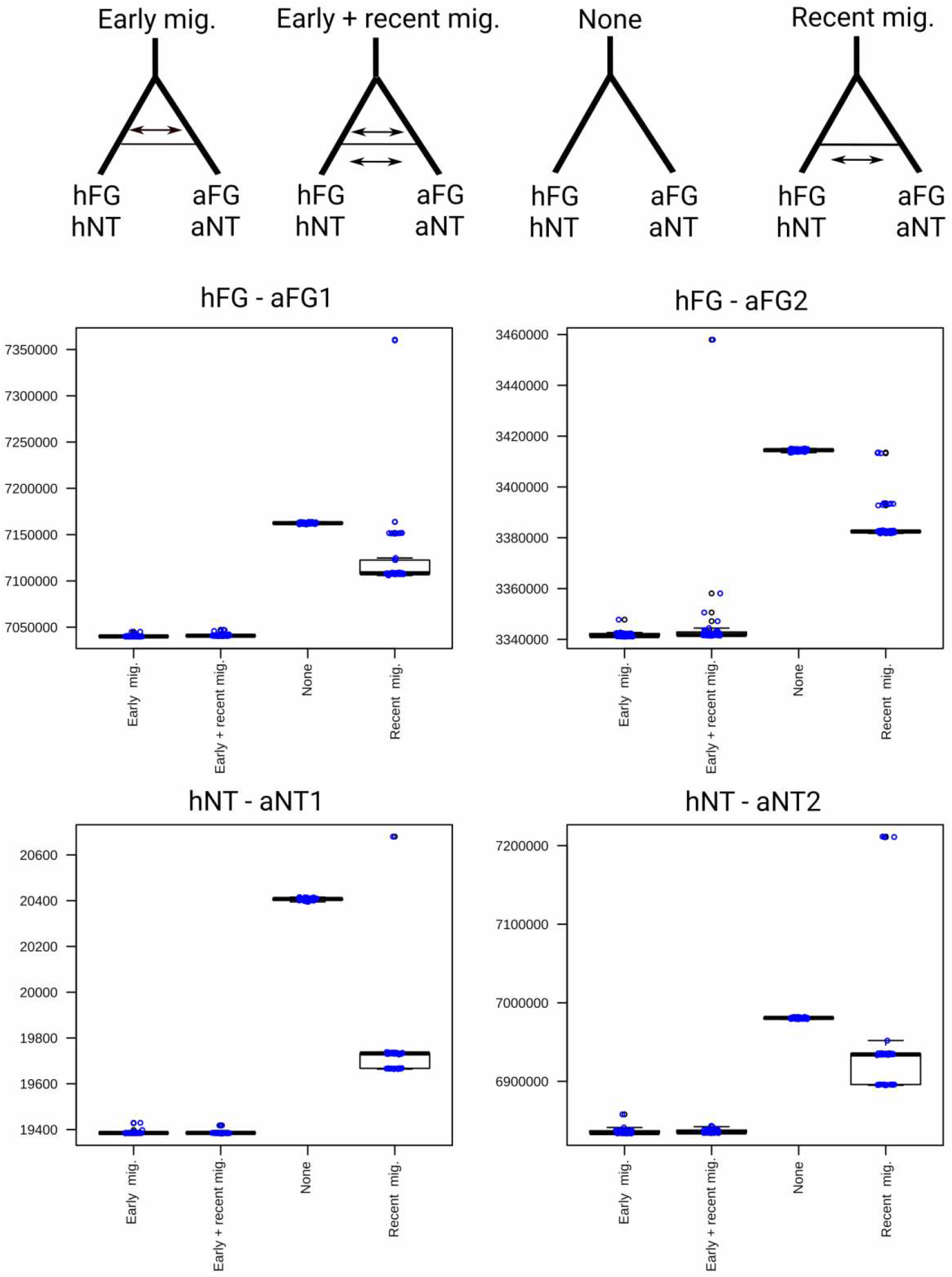
Comparison of Akaike information criteria (AIC) over scenarios of migration and absence of migration between sympatric alpine populations of *Arabidopsis arenosa* (aFG, aNT) and *A. halleri* (hFG, hNT). Model assuming early (i.e. pre-dating the last glacial maximum) migration (Early mig.) was consistently and significantly prioritized over the model assuming ongoing gene flow between alpine populations of both species (Recent mig.) and the absence-of-migration model (None) (p-value < 0.001 in all cases, Tukey HSD test; see upper part of the figure for visualization of models). Adding recent migration to the old migration model (Early + recent mig.) did not significantly increase the model fit (differences in AIC between Early + recent mig. vs Early mig.; p = 0.28 – 0.99, Tukey HSD test). Overall, these results suggest that ongoing gene flow between sympatric alpine populations is not supported by the simulations. Each scenario was simulated by 50 independent fastsimcoal2 runs; the corresponding distribution of AIC values over these 50 runs is shown (blue dots and boxplots).

**Figure S13:**
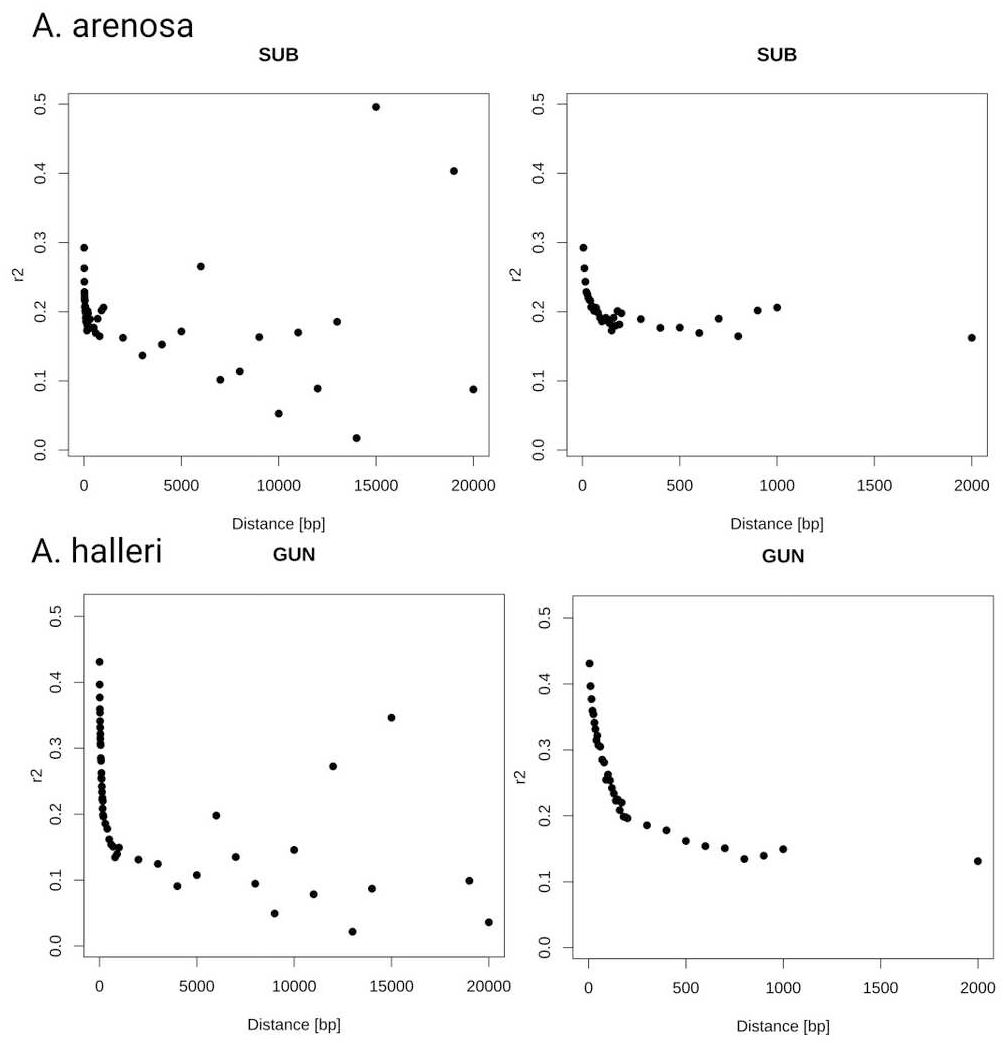
Decay in genotype correlations (r2; a proxy for linkage disequilibrium) over distances along chromosome. Upper line: diploid population with highest coverage of *Arabidopsis arenosa* (SUB of aVT), lower line: diploid population with highest coverage of *A. halleri* (GUN of hNT). The right pane zooms at smaller distances classes.

**Figure S14:**
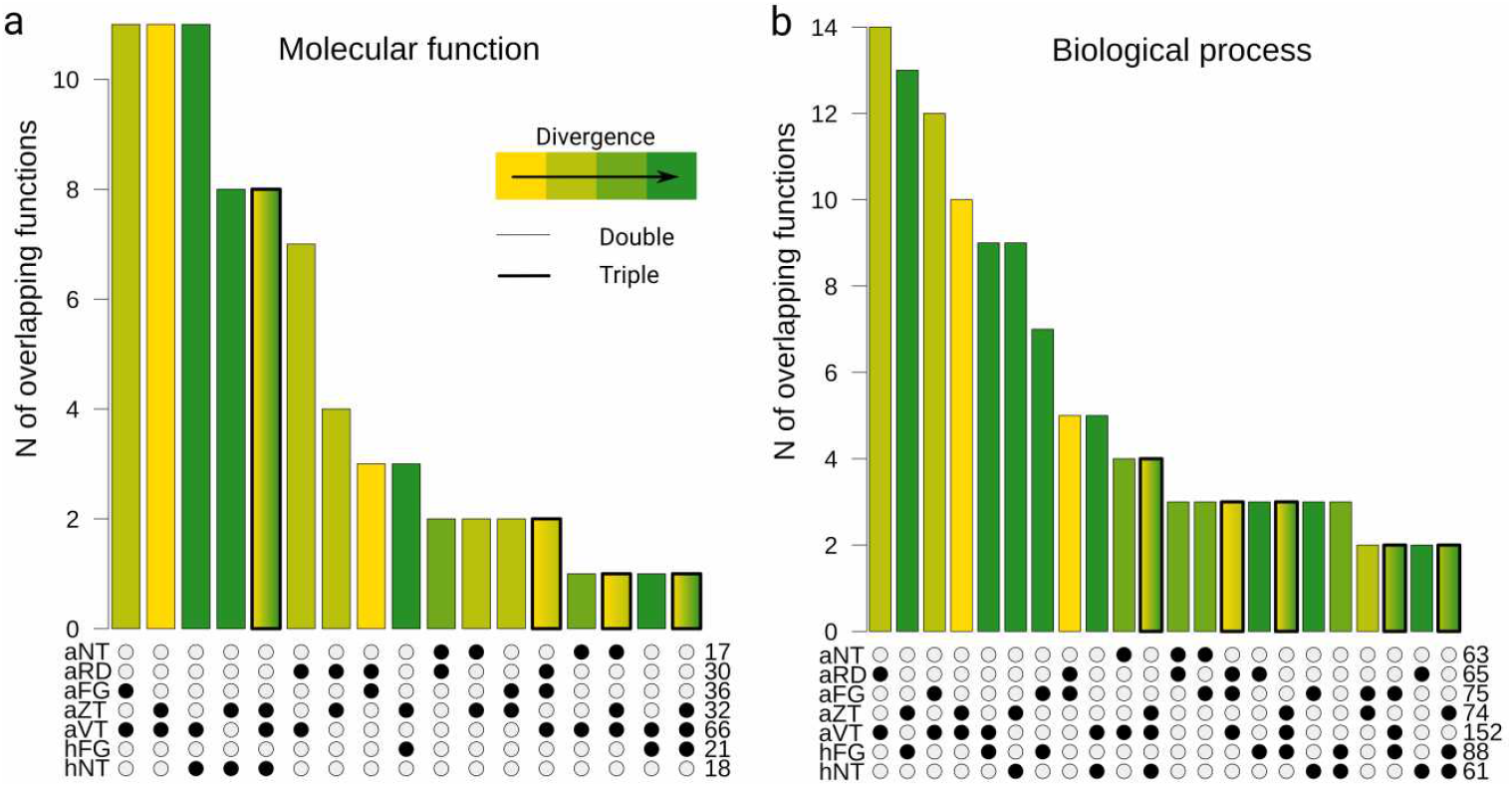
Re-analysis of parallelism by function with ‘molecular function’ ontology. No relationship was identified between divergence and the proportion of parallel candidates by function and this finding was consistent for enriched ‘molecular function’ (a) as well as ‘biological process’ (b, also used in the main text) ontology terms (rM = 0.19 and 0.06, p = 0.3 and 0.6 for molecular function and biological process, respectively, Mantel test of relationship between Jaccard’s similarity in candidate function identity among lineages and 4d-F_ST_). Barplots summarize numbers of parallel GO terms, colored by increasing divergence between parallel lineages in *Arabidopsis arenosa* and *A. halleri*. Only overlaps of > 1 GO term are shown. Numbers in the bottom-right corner of each panel show the total number of candidate items in each region. All displayed categories exhibited parallelism (i.e. significant overlap in the Fisher’s exact test). For region codes see Fig. 1b. Categories with overlap over more than two regions are framed in bold and filled by a gradient.

**Figure S15:**
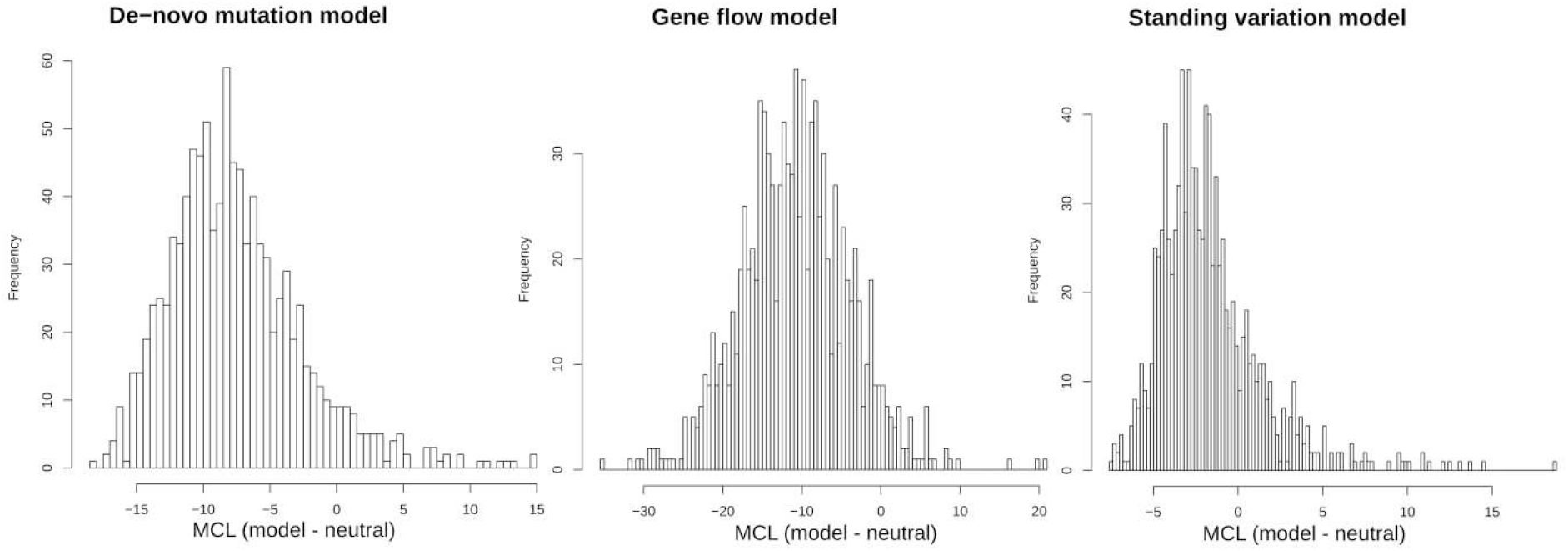
Difference in maximum composite log-likelihoods (MCLs) between parallel and neutral model estimated by DMC for the simulated neutral data (distribution over 1000 simulations). In following analyses, we considered the difference between parallel and neutral model significant if its MCL estimate was higher than the maximum of the distribution of the differences in the simulated data (i.e. above 21).

**Figure S16:**
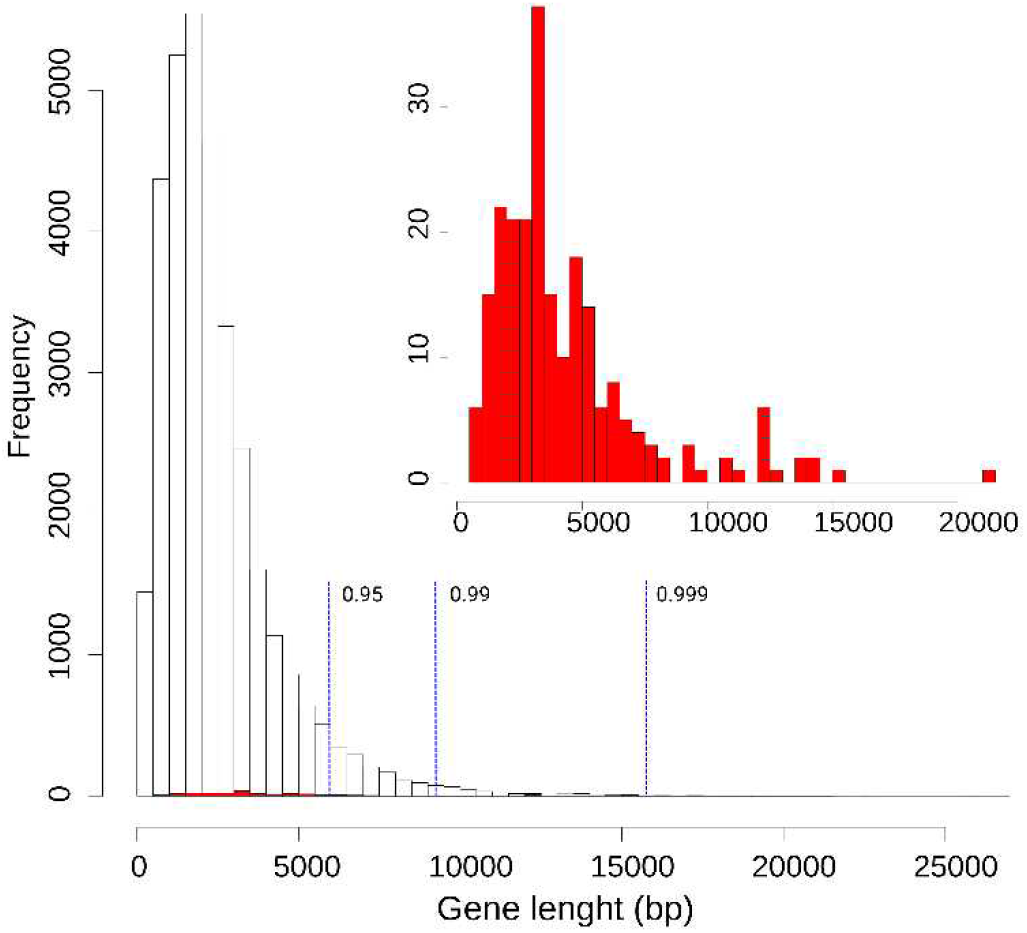
The distribution of genome-wide (white) and parallel gene candidates (red, zoom-in in the inset) gene lengths. Parallel candidate genes are significantly longer than expected given the genome-wide distribution of gene lengths (Table S11), however, only a single parallel gene candidate is an extreme outlier in the gene length. Vertical blue lines show percentiles of distribution of genome-wide gene lengths.

## Supplementary Tables

**Table S1:** List of sampled and sequenced populations of *Arabidopsis arenosa* and *A. halleri.*

| species | lineage | lineage_name | pop | ploidy | A/F | N_indiv | N | E | altitude (m) |
| --- | --- | --- | --- | --- | --- | --- | --- | --- | --- |
| *A. arenosa* | aVT | Vysoké Tatry | VEL | 2x | A | 8 | 49.1620 | 20.1542 | 1823 |
| *A. arenosa* | aVT | Vysoké Tatry | ZEP | 2x | A | 8 | 49.2065 | 20.2151 | 1625 |
| *A. arenosa* | aVT | Vysoké Tatry | SUB | 2x | F | 8 | 48.9603 | 20.3833 | 600 |
| *A. arenosa* | aVT | Vysoké Tatry | BAB | 2x | F | 8 | 49.0435 | 20.1808 | 844 |
| *A. arenosa* | aNT | Niedere Tauern | SCH | 4x | A | 7 | 47.2778 | 14.3214 | 2225 |
| *A. arenosa* | aNT | Niedere Tauern | WIL | 4x | A | 8 | 47.3256 | 14.2304 | 2117 |
| *A. arenosa* | aNT | Niedere Tauern | ING | 4x | F | 8 | 47.2842 | 14.6819 | 970 |
| *A. arenosa* | aNT | Niedere Tauern | KAS | 4x | F | 8 | 46.6883 | 14.8717 | 620 |
| *A. arenosa* | aFG | Făgăraș | LAC | 4x | A | 8 | 45.5954 | 24.6346 | 2092 |
| *A. arenosa* | aFG | Făgăraș | BAL | 4x | A | 8 | 45.6020 | 24.6226 | 2269 |
| *A. arenosa* | aFG | Făgăraș | DRA | 4x | F | 8 | 45.4416 | 25.2239 | 858 |
| *A. arenosa* | aFG | Făgăraș | TIS | 4x | F | 8 | 45.5700 | 25.6083 | 797 |
| *A. arenosa* | aRD | Rodna | INE | 4x | A | 7 | 47.5273 | 24.8806 | 2017 |
| *A. arenosa* | aRD | Rodna | CAR | 4x | F | 8 | 47.5759 | 25.0771 | 981 |
| *A. arenosa* | aZT | Západné Tatry | TKO | 4x | A | 8 | 49.2045 | 19.7352 | 1783 |
| *A. arenosa* | aZT | Západné Tatry | TRT | 4x | A | 8 | 49.2480 | 20.2046 | 1750 |
| *A. arenosa* | aZT | Západné Tatry | HRA | 4x | F | 8 | 49.0072 | 20.2864 | 720 |
| *A. arenosa* | aZT | Západné Tatry | SPI | 4x | F | 8 | 48.9830 | 20.7775 | 552 |
| *A. halleri* | hNT | Niedere Tauern | OBI | 2x | A | 8 | 46.5036 | 14.4870 | 2094 |
| *A. halleri* | hNT | Niedere Tauern | GUN | 2x | F | 8 | 46.9612 | 15.2149 | 612 |
| *A. halleri* | hFG | Făgăraș | HCA | 2x | A | 8 | 45.6009 | 24.6274 | 2251 |
| *A. halleri* | hFG | Făgăraș | DRG | 2x | F | 8 | 46.8636 | 22.8079 | 560 |

Notes: A/F – alpine or foothill population, N\_indiv – number of individuals sequenced per population.

**Table S2:** Number of populations, SNPs per region and neutral differentiation of alpine and foothill populations (F_ST_ A-F) in the final filtered dataset

| lineage | species | N pop | N SNPs | $F_{ST}$ A-F |
| --- | --- | --- | --- | --- |
| aNT | *A. arenosa* | 4 | 11100222 | 0.08 - 0.12 |
| aFG | *A. arenosa* | 4 | 11426015 | 0.10 - 0.18 |
| aRD | *A. arenosa* | 2 | 12053785 | 0.06 |
| aVT | *A. arenosa* | 4 | 11522515 | 0.11 - 0.13 |
| aZT | *A. arenosa* | 4 | 11093985 | 0.04 - 0.07 |
| hNT | *A. halleri* | 2 | 6796677 | 0.24 |
| hFG | *A. halleri* | 2 | 6568335 | 0.19 |
| All *A. arenosa* | *A. arenosa* | 18 | 11351255 | 0.04 - 0.18 |
| All *A. halleri* | *A. halleri* | 4 | 6713051 | 0.19 - 0.24 |

**Table S3:**
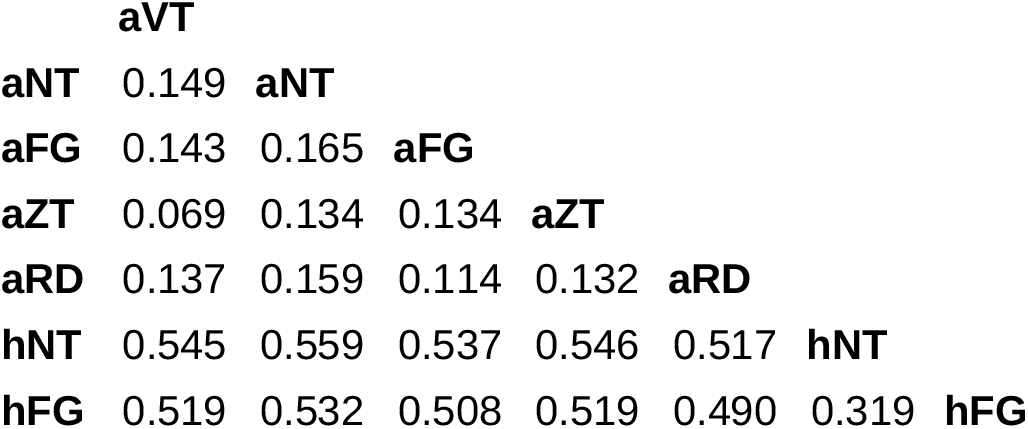
Genome-wide differentiation of the regions. Hudson’s F_ST_ calculated among regions over all 4-fold degenerate sites and averaged over foothill populations (considered ancestral) from each region.

**Table S4:** Per population nucleotide diversity and Tajimas’D calculated over genome-wide 4-fold degenerate sites (subsampled to six individuals per population at each site).

| **lineage** | **pop** | **nucl.diversity** | **Tajimas'D** |
| --- | --- | --- | --- |
| aVT | VEL | 0.023 | 0.08 |
| aVT | ZEP | 0.024 | 0.09 |
| aVT | SUB | 0.027 | -0.03 |
| aVT | BAB | 0.027 | -0.02 |
| aNT | SCH | 0.012 | -0.15 |
| aNT | WIL | 0.027 | 0.36 |
| aNT | ING | 0.025 | 0.48 |
| aNT | KAS | 0.024 | 0.47 |
| aFG | LAC | 0.021 | 0.66 |
| aFG | BAL | 0.026 | 0.41 |
| aFG | DRA | 0.021 | 0.01 |
| aFG | TIS | 0.028 | 0.08 |
| aRD | INE | 0.029 | 0.02 |
| aRD | CAR | 0.021 | 0.11 |
| aZT | TKO | 0.028 | 0.03 |
| aZT | TRT | 0.021 | -0.16 |
| aZT | HRA | 0.027 | 0.01 |
| aZT | SPI | 0.024 | 0.29 |
| hFG | OBI | 0.011 | 0.54 |
| hFG | GUN | 0.015 | 0.50 |
| hNT | HCA | 0.015 | 0.17 |
| hNT | DRG | 0.016 | 0.14 |

**Table S5:** Number of parallel and nonparallel candidate SNPs in the genome (SNPs_full) and particular classes of sites of the genome.

| type | N parallel | N nonparallel | parallel/nonparallel |
| --- | --- | --- | --- |
| SNPs_full | 5615 | 96916 | 0.058 |
| SNPs_genic | 3281 | 44160 | 0.074 |
| SNPs_nonsynonymous | 970 | 13043 | 0.074 |
| SNPs_synonymous | 1286 | 18848 | 0.068 |
| SNPs_upstream | 1844 | 45191 | 0.041 |
| SNPs_downstream | 374 | 3929 | 0.095 |

Notes: The higher proportion of parallel to nonparallel SNPs between SNPs\_full and SNPs\_nonsynonymous is significant ( $p < 0.001$ , Fisher's exact test). Upstream and downstream regions were arbitrarily set at 5 kbs before and after 5' and 3' UTRs

**Table S6:** No evidence that parallel gene candidates are subjected to lower or higher constraints than the rest of the genome.

| lineage | species | N windows genome | N windows parallel | p-value (genome) | p-value (subset) |
| --- | --- | --- | --- | --- | --- |
| aFG | *A. arenosa* | 2741 | 90 | 0.65 | 0.32 |
| aRD | *A. arenosa* | 2713 | 88 | 0.71 | 0.95 |
| aVT | *A. arenosa* | 2572 | 106 | 0.55 | 0.81 |
| aZT | *A. arenosa* | 2728 | 86 | 0.46 | 0.75 |
| aNT | *A. arenosa* | 2060 | 14 | 0.76 | 0.54 |
| hNT | *A. halleri* | 1780 | 20 | 0.75 | 0.66 |
| hFG | *A. halleri* | 2044 | 23 | 0.95 | 0.72 |

Notes: N windows genome: number of 50-kb windows overlapping with any gene in the genome, N windows parallel: number of 50-kb windows overlapping with any parallel gene candidate, p-value: Wilcoxon rank test for difference between $\pi_N/\pi_S$ (used as a proxy for selective constraints) in windows with parallel gene candidates and with any gene genome-wide (p-value genome) or random subset of genome-wide windows equal to the N windows parallel (p-value subset).

**Table S7:** Significant parallelism in alpine adaptation is ubiquitous at gene- and functional-levels in *Arabidopsis arenosa* and *A. halleri.*

| Overlap | % of significant overlaps |  |  |  |
| --- | --- | --- | --- | --- |
|  | by gene |  | by function |  |
|  | all | intraspecific | all | intraspecific |
| two regions | 71% | 91% | 81% | 82% |
| three regions | 54% | 100% | 31% | 30% |
| four regions | 23% | 100% | 3% | 0% |
| five regions | 5% | 100% | 0% | 0% |

Note: Shown are percentages of lineage pairs (21 in total, 11 intraspecific) with significant overlaps between the gene and function candidates ( $p < 0.05$ , Fisher's exact test).
Gene and function counts used to calculate Fisher's exact with corresponding p-values are summarized in Dataset S4 and S5 for candidate gene and function lists, respectively.

**Table S8:**
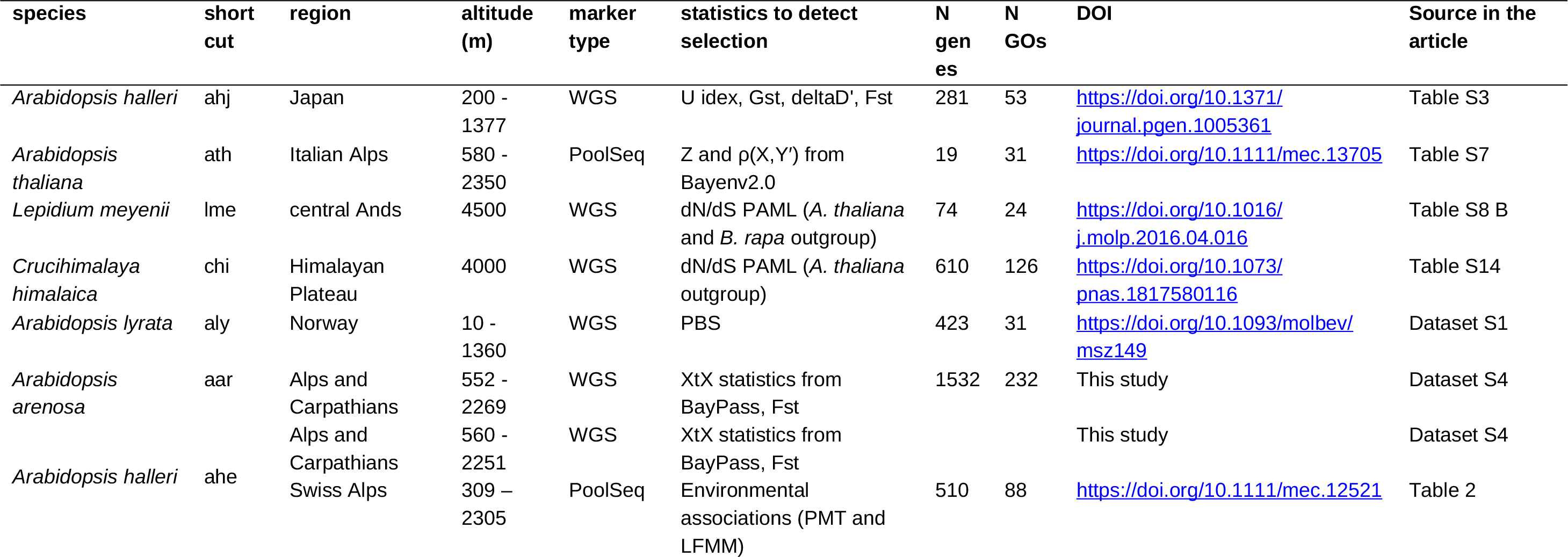
Details on eight genomic studies on alpine adaptation in species from family Brassicaceae used in metaanalysis of gene- and functional-level parallelism.

**Table S9:** Divergence between the species from family Brassicaceae used in our analysis. The values are published divergence time estimates based on the dating of the plastome phylogeny in BEAST (17, 28).

| species 1 | species 2 | divergence (mya) | source |
| --- | --- | --- | --- |
| lme | chi | 18.11 | Hohmann et al., 2015 |
| lme | ath | 18.11 | Hohmann et al., 2015 |
| lme | ahe | 18.11 | Hohmann et al., 2015 |
| lme | ahj | 18.11 | Hohmann et al., 2015 |
| lme | aar | 18.11 | Hohmann et al., 2015 |
| lme | aly | 18.11 | Hohmann et al., 2015 |
| chi | ath | 11.48 | Hohmann et al., 2015 |
| chi | ahe | 11.48 | Hohmann et al., 2015 |
| chi | ahj | 11.48 | Hohmann et al., 2015 |
| chi | aar | 11.48 | Hohmann et al., 2015 |
| chi | aly | 11.48 | Hohmann et al., 2015 |
| ath | ahe | 5.97 | Novikova et al., 2016 |
| ath | ahj | 5.97 | Novikova et al., 2016 |
| ath | aar | 5.97 | Novikova et al., 2016 |
| ath | aly | 5.97 | Novikova et al., 2016 |
| ahe | ahj | 0.56 | Novikova et al., 2016 |
| aly | ahe | 0.56 | Novikova et al., 2016 |
| aly | ahj | 0.56 | Novikova et al., 2016 |
| aar | ahe | 0.6 | Novikova et al., 2016 |
| aar | ahj | 0.6 | Novikova et al., 2016 |
| aar | aly | 0.6 | Novikova et al., 2016 |

Notes: ahj: *Arabidopsis halleri* from Japan, ath: *Arabidopsis thaliana*, lme: *Lepidium meyenii*, chi: *Crucihimalaya himalaica*, aly: *Arabidopsis lyrata*, aar: *Arabidopsis arenosa*, ahe: *Arabidopsis halleri* from Southeastern Alps + Carpathians and *Arabidopsis halleri* from Swiss Alps. See Table S8 for details on studies.

**Table S10:** Parameters used for estimating composite likelihoods of each selection model. In black bold are borderline times used to distinguish between gene-flow/standing variation (lower values) and standing variation/*de-novo* origin (higher values) models. We also used lower (red italics) and higher (green italics) borderline times yet with no qualitatively different results in regards to the relationship between parallelism and divergence (Supplementary Text 5).

| parameter | convergent model | within species | between species |
| --- | --- | --- | --- |
| selection coefficient | all | 0.0001, 0.001, 0.01, 0.05 | 0.0001, 0.001, 0.01, 0.05 |
| migration rates | gene flow | 0.00001, 0.001, 0.1 | 0.00001, 0.001, 0.1 |
| frequencies of standing variant at the onset of selection | standing var. | 6.25e-07, 0.0001, 0.001 | 6.25e-07, 0.0001, 0.001 |
| times for which allele was standing prior to onset of selection | standing var. | 100, **1000**, 5000, 1e4, 5e4, **1e5**, **1e6** | **1e5**, **1e6**, 1.5e6, **2e6**, **2.5e6**, **3e6**, 4e6 |

**Table S11:** Parallel gene candidates are significantly longer than other genes in the genome.

| lineage | N genes genome | N genes parallel | median genome (bp) | median genes parallel (bp) | p-value (genome) | p-value (subset) |
| --- | --- | --- | --- | --- | --- | --- |
| aFG | 33390 | 99 | 2008 | 3401 | 4.00E-16 | 5.33E-07 |
| aRD | 33390 | 103 | 2008 | 3405 | 5.92E-16 | 8.19E-09 |
| aVT | 33390 | 119 | 2008 | 3214 | 2.20E-16 | 1.05E-10 |
| aZT | 33390 | 92 | 2008 | 3568 | 2.20E-16 | 1.58E-12 |
| aNT | 33390 | 22 | 2008 | 4729 | 4.53E-08 | 1.40E-04 |
| hNT | 33390 | 35 | 2008 | 3141 | 2.08E-05 | 5.91E-03 |
| hFG | 33390 | 29 | 2008 | 3456 | 2.26E-06 | 1.53E-04 |

Notes: p-value: Wilcoxon rank test for difference between gene length of parallel gene candidates and all genes genome-wide (p-value genome) or random subset of genome-wide windows equal to the N windows parallel (p-value subset)

## Supplementary Datasets

**Dataset S1:** Case studies of parallel genetic evolution from isolated model systems categorized based on divergence between parallel lineages.

**Dataset S2:** Parallel gene candidates annotated into functions using the TAIR database and associated literature.

**Dataset S3:** Significantly enriched gene ontology terms of parallel gene candidates.

**Dataset S4:** The list of gene candidates within each *Arabidopsis* lineage and their overlap among lineages.

**Dataset S5:** The list of function candidates within each *Arabidopsis* lineage and their overlap among lineages.

**Dataset S6:** The list of gene candidates from each species from the family Brassicaceae and their overlap.

**Dataset S7:** The list of function candidates from each species from the family Brassicaceae and their overlap.

**Dataset S8:** The most likely source of selected variant(s) inferred for each parallel gene candidate.

**Dataset S9:** Sequence processing quality assessment of each re-sequenced individual.

**Dataset S10:**‘Template’ and ‘distribution’ (*tpl* and *est*) files used in coalescent simulations.

